# Tissue-specific biotransformation shapes the toxicity of 6PPD-quinone in fish cell models

**DOI:** 10.64898/2026.11.16.766981

**Authors:** A. Toso, M. E. Franco, T. Masset, W. Dudefoi, P. Nair, S. Goncalves, A. J. Bergmann, E. Vermeirssen, F. Breider, B. J. D. Ferrari, H. Peng, K. Schirmer

## Abstract

Acute mortality of salmonids exposed to urban runoff has been linked to 6PPD-quinone (6PPD-Q), a transformation product of the widely used tire antiozonant 6PPD. However, the cellular processes underlying 6PPD-Q toxicity remain poorly understood. Beyond the well-documented differences in species sensitivity, here we determined tissue-specific responses in rainbow trout (*Oncorhynchus mykiss*) cell lines exposed to 6PPD-Q: brain-derived cells (RTBrain) were more sensitive than gill-derived cells (RTgill-W1). To unravel the cellular mechanisms underlying such differential responses, we combined time-resolved 6PPD-Q cellular depletion kinetics, targeted biotransformation product (BTP) profiling, transporter inhibition experiments, and cytotoxicity assessments. Both cell models biotransformed 6PPD-Q, but RTBrain displayed a 6.6-fold higher cellular clearance rate than RTgill-W1. Chemical profiling identified four BTPs, including *para*-aryl-OH-6PPD-Q, which was selectively toxic to RTBrain but not RTgill-W1 cells. These findings reveal tissue-specific toxicokinetics as potential determinants of differential 6PPD-Q toxicity even within the same species. More broadly, this integrated approach highlights the importance of considering biotransformation and the production of biologically active transformation products when assessing susceptibility to emerging contaminants.

**Teaser:** Biotransformation shapes tissue-specific toxicity of the tire-derived pollutant 6PPD-quinone *in vitro*.

## INTRODUCTION

N-(1,3-dimethylbutyl)-N′-phenyl-p-phenylenediamine (6PPD) is one of the most widely used tire antiozonants worldwide (*1*). Due to its continuous release from tire wear particles, both 6PPD and its transformation product, 6PPD-quinone (6PPD-Q), are frequently detected in urban waters and road runoff (*1–3*). 6PPD-Q was identified as the primary toxicant responsible for urban runoff mortality syndrome (URMS) causing acute mortality in salmonids (*4*, *5*). The concentrations of 6PPD-Q in surface waters have been reported to reach 1.46 µg/L (*6*), exceeding acute toxicity thresholds for sensitive fish species (*4*, *7–9*). Coho salmon (*Oncorhynchus kisutch*) is considered the most sensitive fish identified to date, with reported 24-h LC_50_ values of 0.041–0.095 µg/L (*4*, *7*). Most salmonids, including rainbow trout (*Oncorhynchus mykiss)*, are also susceptible to 6PPD-Q toxicity (*8–10*), but their sensitivity varies substantially among species (*9*, *11*, *12*). Importantly, the lack of a clear phylogenetic pattern suggests that differences in physiological or toxicokinetic mechanisms, rather than evolutionary relatedness alone, may underlie the susceptibility to 6PPD-Q (*8*, *9*, *11*).

Beyond acute mortality, 6PPD-Q has been associated with neurological impairments in exposed fish (*10*, *13*) and adverse effects across multiple tissues (*14*, *15*). Recent evidence has linked 6PPD-Q exposure to oxidative stress, mitochondrial dysfunction, inflammation, and blood– brain barrier disruption (*14*, *16–19*). However, ROS-driven oxidative stress is a well-known consequence of quinone toxicity (*20*, *21*), while mitochondrial respiration is a conserved cellular process (*22*). Therefore, these processes alone cannot explain the observed differences in 6PPD-Q sensitivity (*8*, *11*). Additional toxicokinetic processes, including cellular uptake, transport, biotransformation, and elimination (*e.g.,* via ABCC transporters), as well as the formation of toxic biotransformation products (BTPs), may play a central role in determining sensitivity to 6PPD-Q (*11*, *18*, *23*). These processes can be investigated under controlled *in vitro* conditions using non-animal methods such as fish cell models (*24*, *25*).

Here, we investigated 6PPD-Q and its BTPs in two rainbow trout (RT) cell models representing different tissues: RTgill-W1 and RTBrain, derived from gill and brain, respectively. We observed a greater sensitivity of RTBrain cells to 6PPD-Q compared with that previously observed in RTgill-W1 cells using OECD TG 249-based viability assays (*26*). Such observations led us to hypothesize that toxicokinetic processes, particularly biotransformation pathways, are key determinants of differences in cellular susceptibility. Given the higher expression of several ABCC transporters in RTgill-W1 than in RTBrain cells (*27*), we further hypothesized that ABCC-mediated efflux could contribute to the lower sensitivity of RTgill-W1 to 6PPD-Q. We tested these hypotheses by integrating: i) time-resolved 6PPD-Q depletion kinetics and high-resolution targeted BTP profiling to characterize biotransformation; ii) ABCC inhibition experiments to investigate transporter-mediated elimination; and iii) OECD TG 249-based cytotoxicity assessment of identified BTPs to evaluate their contribution to cellular susceptibility. This integrated approach enabled us to link tissue-specific chemical fate to differences in cellular susceptibility. More broadly, our study highlights the role of chemical toxicokinetics in shaping cellular responses to 6PPD-Q and illustrates the value of fish cell lines for investigating the mechanisms underlying the toxic effects of emerging pollutants.

## RESULTS

### RTBrain cells exhibit greater 6PPD-Q sensitivity and clearance than RTgill-W1 cells

When assessed using an OECD TG 249-based assay (Fig. 1A), RTBrain cells showed markedly greater sensitivity to 6PPD-Q than previously reported for RTgill-W1 under the same exposure conditions (*26*) (Fig. 1B). Metabolic activity was the most sensitive endpoint (EC_50_ = 8.44 µg/L), followed by lysosomal activity (EC_50_ = 14.66 µg/L) and membrane integrity (EC_50_ = 36.48 µg/L) (Fig. 1B). To characterize 6PPD-Q kinetics in RTBrain and RTgill-W1 cells, we then employed a recently developed cell-based system (*28*) to measure its distribution across the cell culture medium, cell monolayer, and culture vessel surface over time (Fig. 1C). We observed distinct depletion kinetics between the two cell lines over a 48-h exposure period (Figs. 1D-E). In the RTBrain cell system, the 6PPD-Q concentration in the medium rapidly declined by 59% within 4 h and was no longer detectable at 24 h; an additional 2-h sampling point was included for these cells to further resolve the initial depletion phase, with 6PPD-Q showing a 44% decline within 2 h of exposure (Fig. 1D). In contrast, in the RTgill-W1 system, the 6PPD-Q concentration in the medium declined by 25% within 4 h and by 86% at 48 h (Fig. 1E). Extended exposure experiments (96 h) further showed that 6PPD-Q remained detectable in the RTgill-W1 medium throughout the full exposure period, showing a 92% reduction by 96 h (Supplementary Fig. S3). Importantly, mass balance analyses showed only transient cellular association of 6PPD-Q in both cell lines (Fig. 1D-E, Supplementary Fig. S3A-B). Indeed, 6PPD-Q was detected in the cellular fraction at early time points but did not accumulate over time. Most of the recovered 6PPD-Q remained in the exposure medium throughout the experiment, while a small fraction partitioned to the culture vessel, consistent with previous reports on 6PPD-Q stability and its associations with plastic materials (*29*). Cell-free controls demonstrated negligible abiotic loss of 6PPD-Q over the experimental periods, supporting the attribution of 6PPD-Q depletion in cell-containing samples to cell-associated processes (Figs. 1F and Supplementary Figs. S3C-D).

**Figure 1.**
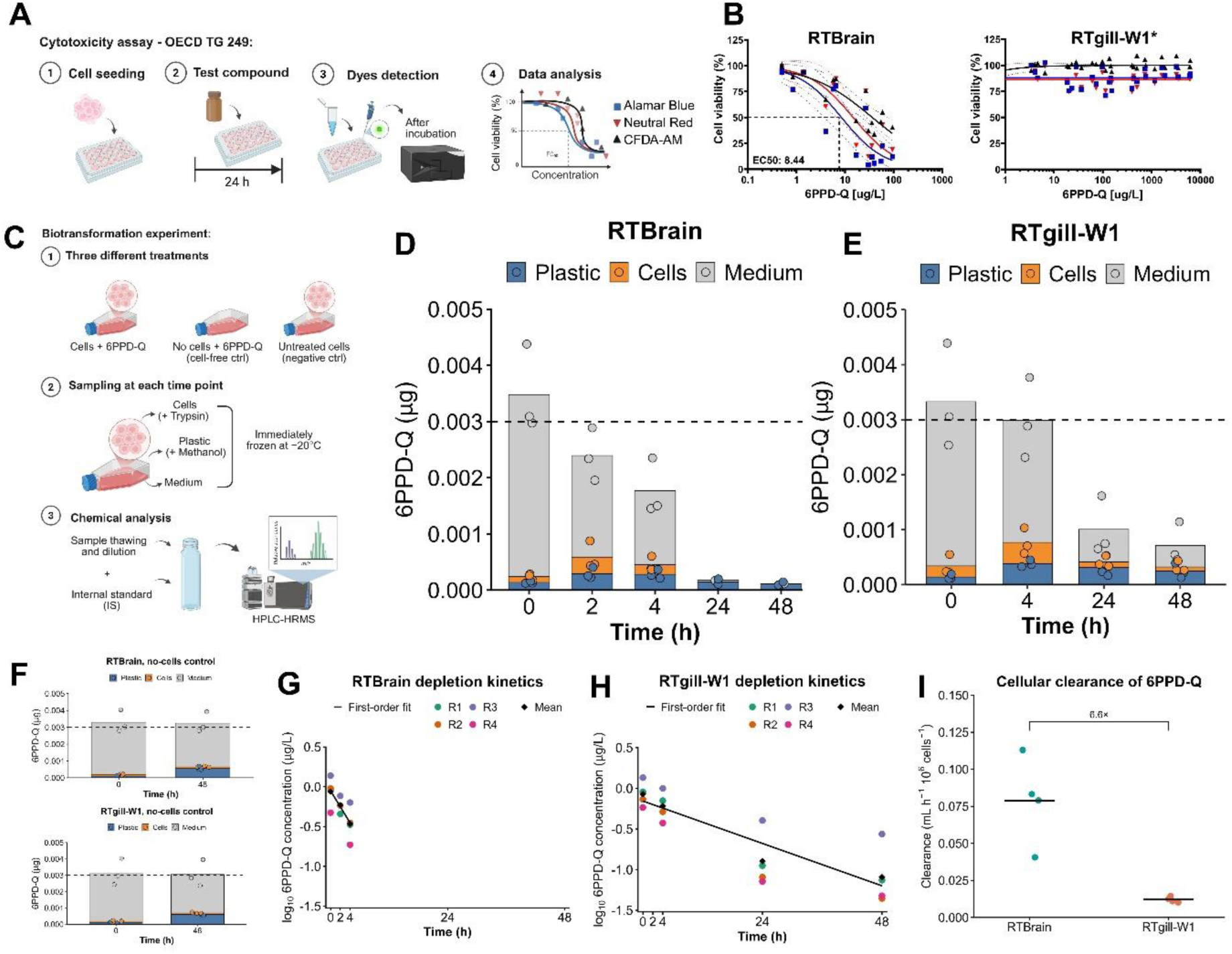
Differential 6PPD-Q sensitivity and toxicokinetics in RTBrain and RTgill-W1 cells. (A) OECD TG 249 workflow. (B) Acute cytotoxicity of 6PPD-Q in RTgill-W1 and RTBrain cells after 24 h of exposure. Data are expressed as percentages of the solvent control (0.5% (v/v) DMSO). Dose-response curves are shown for metabolic activity (AlamarBlue™, blue squares), membrane integrity (CFDA-AM, black triangles), and lysosomal integrity (Neutral Red, red inverted triangles). Dashed lines indicate the 95% confidence bands around the fitted curves, where calculable. Data points represent measured concentrations from individual biological replicates (n = 3). Asterisks indicate previously published data reproduced from Dudefoi et al. (2024) under the terms of the Creative Commons Attribution 4.0 International licence (CC BY 4.0) (*26*). (C) Biotransformation experiment workflow. (D-E) Time-dependent distribution of 6PPD-Q (µg) across medium, cellular, and plastic-associated compartments in RTBrain (D) and RTgill-W1 (E) exposure systems. The dashed line indicates the nominal initial mass of 6PPD-Q (0.003 µg), corresponding to the target exposure concentration of 1 µg/L in the 3-mL exposure volume. (F) Distribution of 6PPD-Q in no-cell controls for RTBrain (upper panel) and RTgill-W1 (lower panel) exposure systems. The same fractionation procedure was applied as for cell-containing cultures to facilitate direct comparison. Consequently, 6PPD-Q detected in this fraction in the absence of cells may reflect residual exposure medium. (G-H) Depletion kinetics of 6PPD-Q from the exposure medium in RTBrain (G) and RTgill-W1 (H). (I) Apparent cellular clearance of 6PPD-Q in RTBrain and RTgill-W1 cells. Data in (D–F) represent three independent biological replicates (n = 3) over 48 h. Data in (G–I) represent individual biological replicates (n = 4), comprising the three biological replicates from the main experiments (whose mass balance is shown in panels D–E) and one additional replicate (R4; Supplementary Fig. S1). For consistency, data from R4 were included only up to 48 h, although the original experiment extended to 96 h.

Furthermore, kinetic analyses showed that 6PPD-Q depletion followed first-order kinetics in both cell lines and a faster depletion of the parent compound in the RTBrain system (Figs. 1G-H). The apparent *in vitro* clearance (CL_IN VITRO_) was approximately 6.6-fold higher in RTBrain (0.0789 mL h⁻¹ 10⁶ cells⁻¹) than in RTgill-W1 (0.0120 mL h⁻¹ 10⁶ cells⁻¹), and this difference was statistically significant (Welch’s t-test; p value = 0.020) (Fig. 1I).

### RTBrain and RTgill-W1 display distinct 6PPD-Q biotransformation profiles

We performed a suspect screening analysis using a literature-derived list of reported 6PPD-Q BTPs (Supplementary Table S1). Of the 29 screened compounds, we detected four in the experimental samples (Supplementary Fig. S4); three corresponded to monohydroxylated phase I BTPs (m/z 315.1703), whereas the fourth corresponded to a glucuronide-conjugated monohydroxylated phase II BTP (TP-O-Gluc; m/z = 491.20; RT: 19.7). The three phase I BTPs were unambiguously identified as C4-alkyl-OH-6PPD-Q, C3-alkyl-OH-6PPD-Q, and *para*-aryl-OH-6PPD-Q, with assignment confirmed against synthetic standards (Supplementary Table S2; Supplementary Fig. S5) and further supported by MS/MS fragmentation patterns (Supplementary Figs. S6–S8), consistent with previous reports (*30*, *31*). In contrast, TP-O-Gluc was tentatively identified as a glucuronide-conjugated monohydroxylated 6PPD-Q BTP (m/z 491.20), as no synthetic standard was available to confirm its identity.

The detected BTPs showed distinct temporal profiles between the two cell lines (Supplementary Figs. S9-10). C4-alkyl-OH-6PPD-Q was the predominant phase I BTP in both cell models. In RTBrain medium, this compound peaked early (∼0.2 µg/L within 4 h) before declining below 0.1 µg/L at 24 h (Supplementary Fig. S9A). In the RTgill-W1 medium, it increased progressively to a 24-48 h plateau of ∼0.15 µg/L (Supplementary Fig. S10A). Intracellularly, C4-alkyl-OH-6PPD-Q remained relatively low (<0.1 µg/L) in both cell lines but was exclusively detected at 0–4 h in RTBrain (Supplementary Fig. S9B) and at 24–48 h in RTgill-W1 (Supplementary Fig. S10B). The other phase I BTPs were detected at much lower levels and predominantly in the medium compartment. The C3-alkyl-OH-6PPD-Q appeared only in the RTgill-W1 system, and only at 24– 48 h (<0.1 µg/L; Supplementary Fig. S10C). The *para*-aryl-OH-6PPD-Q was restricted to 0–4 h in RTBrain (<0.1 µg/L; Supplementary Fig. S9C) and exclusively at 24 h in RTgill-W1 (<0.1 µg/L; Supplementary Fig. S10D). Additionally, the tentatively identified phase II metabolite TP-O-Gluc was detected only at late time points in both cell lines, with higher relative abundance in RTgill-W1 than in RTBrain; in RTgill-W1, mean area ratios (compound/IS) were ∼0.03 at 24 h and ∼0.05 at 48 h (Supplementary Fig. S10E) compared with values below 0.02 at both time points in RTBrain (Supplementary Fig. S9D).

### *Para*-aryl-OH-6PPD-Q is the only cytotoxic biotransformation product

The acute cytotoxicity of five synthetic standards representing phase I BTPs of 6PPD-Q was screened using an OECD TG 249-based assay. At a nominal concentration of 100 µg/L, only *para*-aryl-OH-6PPD-Q reduced metabolic activity in RTBrain cells to approximately 45% of control levels, whereas no relevant effects were observed for this compound in RTgill-W1 cells or for the other tested compounds in either cell line (Fig. 2A). Based on this preliminary screening, *para*-aryl-OH-6PPD-Q was further evaluated using a full concentration–response assessment (Fig. 2B-C). In RTBrain cells, metabolic activity was the most sensitive endpoint, with an EC_50_ value of 25.62 µg/L (95% CI: 17.41–37.72 µg/L), followed by lysosomal activity (EC_50_ = 52.13 µg/L; 95% CI: 37.0–73.45 µg/L) and membrane integrity (EC_50_ = 68.92 µg/L; 95% CI: 49.0–96.65 µg/L). These values were calculated using the geometric mean of the measured concentrations at t_0_ and t_24_, corresponding to the beginning and end of the exposure period, respectively (Fig. 2B). EC_50_ values were of the same order of magnitude as those previously reported for the parent compound 6PPD-Q (Fig. 1B). No consistent cytotoxicity was observed across biological replicates in RTgill-W1 cells (Fig. 2C). This result was in agreement with previously reported tissue-specific responses to the parent compound under the same exposure conditions (*26*). Interestingly, *para*-aryl-OH-6PPD-Q concentrations decreased by 32.6% in RTBrain medium and by 62.1% in RTgill-W1 medium over the 24-h exposure period (Supplementary Table S3). Microscopic inspection revealed clear morphological changes in RTBrain cells exposed to *para*-aryl-OH-6PPD-Q, including cell rounding and apparent detachment, accompanied by membrane blebbing and an increase in cell-free areas (Fig. 2D), consistent with cellular injury and cytotoxicity (*32*). No morphological changes were observed in RTgill-W1 cells (Fig. 2D).

**Figure 2.**
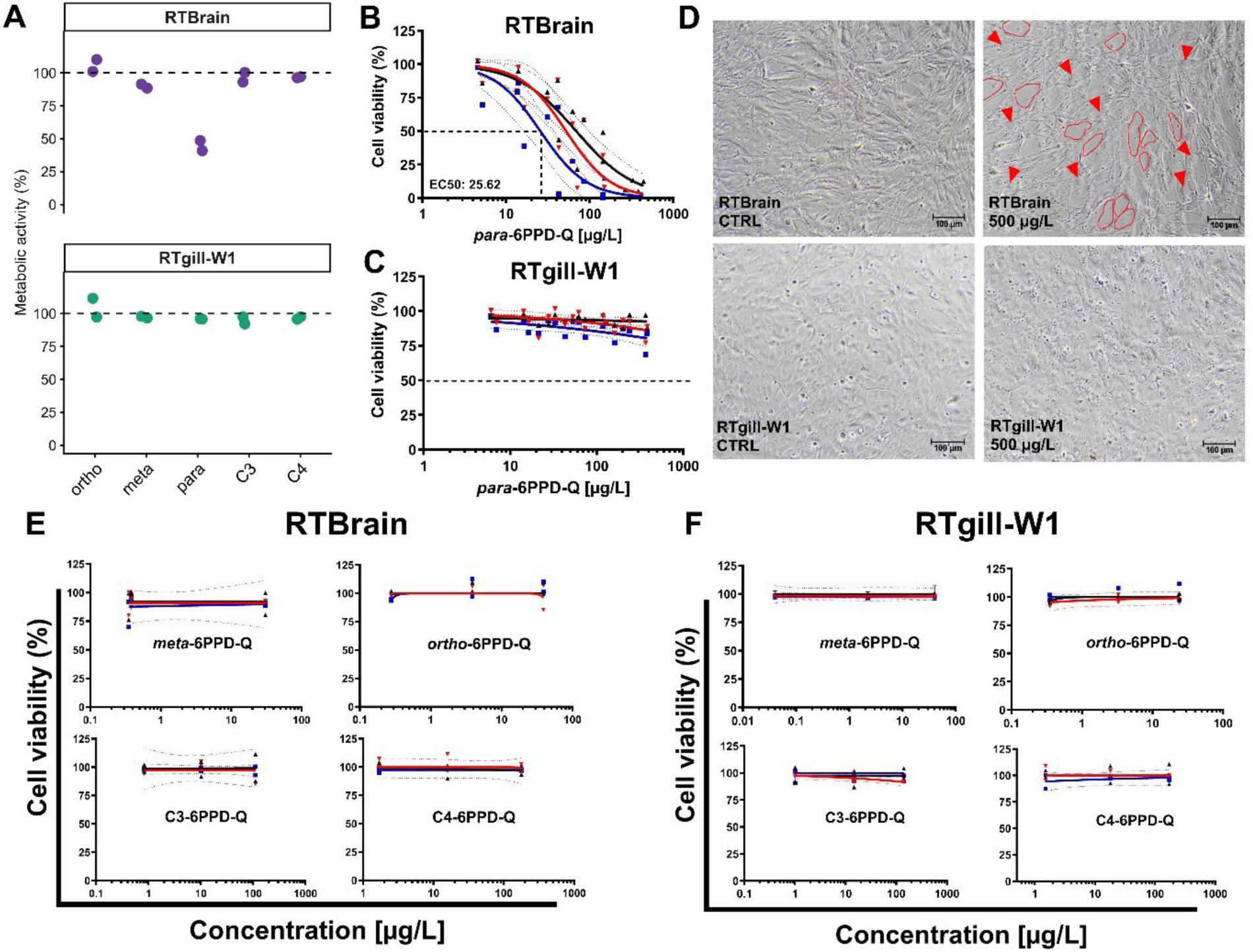
Acute cytotoxicity screening of 6PPD-Q BTPs in RTBrain and RTgill-W1. All data are expressed as percentage of the solvent control (0.5% (v/v) DMSO) with dose-response curves representing respectively cell metabolic activity (AlamarBlue™; blue squares), cell membrane integrity (CFDA-AM; black triangles), and lysosomal integrity (Neutral Red; red inverted triangles) after 24 h of exposure. Dashed lines indicate the 95% confidence bands around the fitted curves, where calculable. (A) Comparative screening of phase I 6PPD-Q BTPs at 100 µg/L nominal concentration. Data represent two preliminary biological replicates based on AlamarBlue™ metabolic activity. (B-C) Full dose–response assessment of *para*-aryl-OH-6PPD-Q in RTBrain (B) and RTgill-W1 cells (C) based on measured concentrations. Data are presented as individual biological replicates (n = 3). SDS positive control was included in all biological replicates (Supplementary Table S5 and Fig. S16). (D) Representative bright-field microscopy images of RTBrain and RTgill-W1 cells in both solvent controls (CTRL) and cells exposed to *para*-aryl-6PPD-Q (500 µg/L); arrows indicate rounded cells, while outlined areas delineate cell-free regions. (E-F) Dose– response curves of four BTPs in RTBrain (E) and RTgill-W1 (F) cells. No concentration-dependent cytotoxicity was observed within the tested concentration range. Data are presented as individual biological replicates (n = 2); analytical verification of exposure concentrations and SDS positive control were performed in one biological replicate. The resulting exposure concentrations were considered representative of the exposure conditions applied in both biological replicates.

The lack of observed cytotoxicity of the other BTPs, including alkyl-hydroxylated isomers (C3- and C4-alkyl-OH-6PPD-Q) and aryl-hydroxylated isomers (*meta*- and *ortho*-aryl-OH-6PPD-Q) (Fig. 2E-F) further highlights the selective toxicity of *para*-aryl-OH-6PPD-Q among the tested BTPs. Interestingly, chemical analysis of the exposure media collected during the BTP cytotoxicity experiments showed that TP-O-Gluc was detected in RTgill-W1 but not RTBrain medium following exposure to *para*-aryl-OH-6PPD-Q (Supplementary Fig. S11).

Lastly, additional experiments using six structurally related PPD-quinones (IPPD-Q, CPPD-Q, HPPD-Q, OPPD-Q, DPPD-Q, and DTPD-Q) (Supplementary Figs. S12-13) revealed no detectable cytotoxicity in either cell line, further suggesting that structural features specific to 6PPD-Q may contribute to its cytotoxicity.

### ABCC inhibition did not increase cytotoxicity in RTgill-W1

Based on findings from our previous study reporting differences in ABCC transporter expression between the two cell lines (*27*), with higher gene expression of several ABCC transporter in RTgill-W1 than in RTBrain cells, we investigated whether ABCC-mediated efflux contributes to the lower sensitivity of RTgill-W1 cells to 6PPD-Q (*26*). Co-exposure to 6PPD-Q and the ABCC inhibitor MK-571 (2 µM) during cell viability experiments (further methodological details are provided in Supplementary Text S4.2) showed that ABCC inhibition did not significantly alter 6PPD-Q-induced cytotoxicity under the tested condition (Supplementary Fig. S14). This lack of acute toxicity suggests that ABCC-mediated efflux is unlikely to account for the resistance of RTgill-W1 cells to 6PPD-Q under these experimental conditions.

## DISCUSSION

Species-specific responses to 6PPD-Q exposure have revealed differences in susceptibility among fish species, with rainbow trout being highly susceptible among teleost fish (LC_50_ = 0.90-1.96 µg/L) (*9*, *10*). Previous *in vivo* studies have implicated toxicokinetic processes, including biotransformation, as key determinants of this species-specific susceptibility (*11*, *30*, *33*). Our findings extend this knowledge to differences in cellular sensitivity and biotransformation in cells derived from tissues of the same species. Specifically, we showed that 6PPD-Q toxicity can differ substantially between gill- and brain-derived cells. By integrating tissue-specific toxicokinetics, transport-mediated elimination, and intrinsic cellular susceptibility in RTgill-W1 and RTBrain cells, our study provides important mechanistic insight into these cellular differences, further highlighting the relevance of brain-specific responses considering the potential neurotoxic effects of 6PPD-Q (*10*, *13*, *17*, *34*, *35*).

### Tissue-specific biotransformation shapes 6PPD-Q cellular fate

Biotransformation processes are a key component of an organism’s defense against harmful chemicals, potentially influencing its tolerance or susceptibility to toxic compounds (*36*, *37*). Although the liver and, to a lesser extent, the gills are commonly investigated sites of xenobiotic biotransformation in fish (*25*, *38*, *39*), the biotransformation capacity of the brain may warrant greater consideration (*25*). Evidence from human toxicology indicates that both brain tissue and the blood–brain barrier are metabolically active sites which can contribute to xenobiotic biotransformation (*40–42*). This local metabolic capacity may thus represent an important, yet comparatively understudied, component of neurotoxicological responses.

Importantly, our results clearly demonstrate that both RTgill-W1 and RTBrain cells can biotransform 6PPD-Q, although with cellular clearance values (0.0120 and 0.0789 mL h⁻¹ 10⁶ cells⁻¹, respectively) lower than the value reported for the liver-derived rainbow trout cell line RTL-W1 (9.88 mL h⁻¹ 10⁶ cells⁻¹) (*43*), consistent with the generally reduced biotransformation capacity of extrahepatic tissues (*24*, *25*). Despite the greater sensitivity of RTBrain to 6PPD-Q, the clearance in these cells was approximately 6.6-fold higher than in RTgill-W1 cells. This differential response raises the question of whether biotransformation primarily contributes to detoxification of 6PPD-Q or rather generates active products which may overall contribute to its toxicity burden.

When resolving the BTP profile, we detected multiple phase I BTPs in both cell types, consistent with findings from previous *in vitro* and *in vivo* studies (*18*, *23*, *30*, *31*, *33*, *43*, *44*). Although both cell lines generated the same major phase I BTPs, their formation kinetics differed. C4-alkyl-OH-6PPD-Q, the most abundant BTP detected in both cell lines, peaked earlier in RTBrain medium and then rapidly declined, while it levels increased over time and reached a plateau in RTgill-W1 medium (Supplementary Figs. S9-10). Interestingly, *para*-aryl-OH-6PPD-Q was detected only at early time points in RTBrain medium and only at 24 h in RTgill-W1 medium, while C3-alkyl-OH-6PPD-Q was only detected in RTgill-W1 medium at low concentrations (Supplementary Figs. S9-10).

The assessment of the acute toxicity of these three phase I BTPs revealed that only *para*-aryl-OH-6PPD-Q induced cytotoxic effects in RTBrain cells, whereas no effects were observed in RTgill-W1 cells (Fig. 2A-B). This pattern mirrors the greater sensitivity of RTBrain to the parent compound 6PPD-Q. Neither C3-nor C4-alkyl-OH-6PPD-Q induced cytotoxicity in either cell line (Figs. 2E-F). We also observed that the position of the hydroxyl group on the aromatic ring seems to strongly influence cytotoxicity, as neither the *ortho*-nor the *meta*-isomer was toxic in RTBrain cells (Figs. 2E-F). A similar regioselective toxicity pattern was recently reported by Nair et al. (2025) (*31*) in coho salmon embryo cells (CSE-119), where aryl-OH-6PPD-Q isomers ranked in toxicity as *para* > *meta* > *ortho*.

It has also been suggested that 6PPD-Q hydroxylation, mediated by CYP450 enzymes, is a primary route of 6PPD-Q detoxification both in humans and in fish (*45*, *46*). Tissue-specific differences in the expression and activity of biotransformation enzymes (*24*, *25*) may therefore contribute to differences in 6PPD-Q biotransformation between RTBrain and RTgill-W1. However, identifying the specific CYP450 isoforms responsible for *para*-aryl-OH-6PPD-Q formation remains challenging, as fish CYP450 enzymes differ substantially from their mammalian counterparts, limiting the applicability of available isoform-selective inhibitors commonly used in human toxicology studies (*30*, *47*).

Furthermore, we also (tentatively) identified the phase II metabolite TP-O-Gluc in the biotransformation experiments in both cell lines. Its detection, previously reported both *in vitro* (*43*), and *in vivo* (*11*, *23*, *33*), as well as in human biological fluids (*45*), indicates that 6PPD-Q undergoes glucuronidation, which generally increases a compound’s hydrophilicity thereby facilitating its detoxification (*45*). Interestingly, we observed distinct formation kinetics in both cell lines as TP-O-Gluc relative abundance remained quite stable at very low levels in RTBrain medium while it considerably increased over time in RTgill-W1 medium (Supplementary Figs. S9-10). These contrasting temporal profiles of TP-O-Gluc in the two cell lines may reflect differences in the formation or accumulation of this glucuronidated BTP. Moreover, measurements during the cytotoxicity assays with *para*-aryl-OH-6PPD-Q showed that TP-O-Gluc was detected only in RTgill-W1 but not in RTBrain (Supplementary Fig. S11), while *para*-aryl-OH-6PPD-Q concentrations decreased by 32.6% in RTBrain medium and 62.1% in RTgill-W1 medium (Supplementary Table S3).

Overall, our results suggest that RTBrain cells biotransform 6PPD-Q more rapidly, while RTgill-W1 cells show evidence of an additional downstream processing of *para*-aryl-OH-6PPD-Q. This pattern raises the possibility that differences in phase II biotransformation between the two cell lines contribute to the distinct cellular responses to 6PPD-Q. In this context, an imbalance between phase I BTP formation and subsequent phase II conjugation has been proposed *in vivo* as a metabolic “bottleneck” that could contribute to the persistence of unconjugated metabolites and increased toxicity (*33*). However, the phase I BTPs giving rise to TP-O-Gluc in the biotransformation experiments could not be directly established. Their observed formation kinetics may also be influenced by rapid turnover, the formation of other phase II BTPs, or the involvement of additional detoxification pathways (*48*, *49*). Moreover, as no synthetic standard was available, TP-O-Gluc could only be tentatively identified. The availability of a synthetic standard would allow its identity to be confirmed and its formation kinetics to be quantified more reliably, providing further insight into its role in 6PPD-Q biotransformation.

### Additional factors contributing to differential 6PPD-Q sensitivity beyond biotransformation

Efflux transporters, including ABC proteins like ABCC transporters, contribute to cellular handling of and defense against xenobiotics by exporting chemicals and their transformation products, hence limiting intracellular exposure (*36*, *50*). Because differences in transporter expression could contribute to tissue-specific 6PPD-Q efflux, we investigated whether ABCC-mediated transport influenced cellular sensitivity to 6PPD-Q. ABCC transporters were of particular interest because their expression has previously been characterized in RTgill-W1 and in RTBrain cells, with overall higher expression reported in RTgill-W1 (*27*). Interestingly, we demonstrated that ABCC inhibition did not increase the sensitivity of RTgill-W1 cells exposed to 6PPD-Q, suggesting that ABCC-mediated efflux is unlikely to be a major determinant of the differential sensitivity observed between the two cell lines.

Beyond transporter-mediated effects, toxicodynamic mechanisms may also contribute to differences in cellular sensitivity to 6PPD-Q. These may involve differences in intrinsic susceptibility to oxidative stress (*16*, *51*) or interactions with specific molecular targets, such as peroxisome proliferator-activated receptor gamma (PPARγ), retinoic acid receptor α (RARα), and retinoid X receptor α (RXRα), which have been suggested as potential 6PPD-Q targets (*16*, *52*, *53*). Given the roles of RARα and RXRα in neural development and function (*54*, *55*), and of PPARγ in regulating metabolism, inflammatory responses, and redox homeostasis in the brain (*56*), these receptors could mediate effects of 6PPD-Q in brain-derived cells, although their contribution to the greater sensitivity of RTBrain remains to be investigated. Oxidative stress may also contribute to the higher sensitivity of RTBrain cells, given the reported susceptibility of brain cells to oxidative imbalance (*57–59*). The quinoid structure of 6PPD-Q and *para*-aryl-OH-6PPD-Q provides a plausible chemical basis for redox perturbation, as quinone-derived electrophiles are known to react with cellular nucleophiles and undergo redox cycling (*21*, *60*, *61*). Differences in the antioxidant defense capacity of RTbrain and RTgill-W1 cells, potentially reflecting differences in their antioxidant enzyme repertoire, may also influence their susceptibility to quinone-induced oxidative stress. Interestingly, redox-cycling potential depends on ring substituents (*21*), suggesting that structural differences among BTPs may influence their potential to induce redox perturbations.

Additionally, 6PPD-Q has been shown to disrupt mitochondrial respiration by uncoupling the electron transport chain (*18*), and to alter energy metabolism in primary dopaminergic neurons (*17*). The reduced metabolic activity observed in the Alamar Blue assay may be consistent with these alterations in (general) cellular energy metabolism. Given the high energy demands of neuronal function and the reliance of brain cells on mitochondrial oxidative phosphorylation to meet these demands (*62*, *63*), RTBrain cells may be particularly vulnerable to mitochondrial dysfunction. However, this hypothesis requires further investigation using direct measurements of mitochondrial function in RTBrain cells, such as oxygen consumption rate and ATP production.

### Implications and perspectives

Despite our important findings on cell-line-specific susceptibility to 6PPD-Q, two key questions remain unresolved. First, to what extent does bioactivation of 6PPD-Q into potentially toxic BTP contribute to the greater toxicity observed in RTbrain cells? The higher clearance observed in these cells, despite their greater susceptibility, suggests that more rapid biotransformation does not necessarily translate into greater cellular protection. Second, although *para*-aryl-OH-6PPD-Q was the only BTP that elicited cytotoxicity in RTBrain cells, whether it is *the ultimate* toxicant of 6PPD-Q remains to be established. Its EC_50_ (25.62 µg/L), which is higher than that of 6PPD-Q (8.44 µg/L), suggests that *para*-aryl-OH-6PPD-Q may account for only part of the toxicity associated with the parent compound, while other (as-yet-unidentified) BTPs may also contribute to the overall toxicity burden.

An important limitation of our study is that, although the biotransformation capacity and toxicokinetic properties of RTBrain cells have already been investigated (*24*, *27*), their cellular identity remains incompletely characterized. However, ongoing transcriptomic analyses by our research group indicate that this cell line comprises a heterogeneous neural-like population dominated by a radial glial-like signature, thus RTBrain cells appear to retain neural stem cell-like, multipotent characteristics, in line with the life-long neurogenic properties of the teleost brain (*64*, *65*). Among rainbow trout cell lines evaluated to date, RTBrain is the only cell line reported to be highly sensitive to 6PPD-Q, whereas other commonly used rainbow trout cell lines, including RTG-2, RTgill-W1, and RTL-W1, have shown lower sensitivity under similar test conditions (*18*, *26*, *66*).

We chose rainbow trout because of its ecological relevance as a cold-water salmonid, its widespread use in aquaculture and aquatic ecotoxicology (*67*), and its documented sensitivity to 6PPD-Q (*9*, *10*). Nonetheless, coho salmon is also highly sensitive to 6PPD-Q (*7*, *19*, *68*), and the coho salmon embryonic cell line, CSE-119, has been reported to exhibit high sensitivity *in vitro* (*30*, *44*, *66*). However, its cellular composition also remains poorly characterized. Notably, previous studies have reported stem-like and progenitor cell characteristics in different fish-derived cell lines, including rainbow trout intestinal cell lines and the liver-derived cell line RTL-W1 (*69*, *70*), as well as an embryo-derived cell line from Atlantic salmon (*Salmo salar*) (*71*). Integrating biotransformation assessment with transcriptomic approaches in both CSE-119 and RTBrain cells could help determine whether the mechanisms identified in this study are shared across these two cellular models and potentially between the two species. Furthermore, other high-resolution approaches, such as single-cell RNA sequencing, could help determine whether the observed responses are specific to brain-derived cells or instead reflect cellular characteristics shared by both cell systems, potentially including stem-like cell populations.

More broadly, the widespread occurrence of 6PPD-Q in aquatic environments (*1*, *72–74*) underscores the need for improved hazard assessment of tire-derived chemicals. In this context, rainbow trout-derived cell models may support the assessment of potential safer alternatives to 6PPD in line with the principles of Safe and Sustainable by Design (SSbD). This is particularly relevant for RTBrain, which showed EC_50_ values within the same order of magnitude as the LC_50_ reported for rainbow trout (0.90-1.96 µg/L) (*9*, *10*). Our findings also demonstrate that both RTBrain and RTgill-W1 are metabolically competent to biotransform 6PPD-Q, consistent with *in vivo* studies (*11*, *30*, *31*). Overall, these findings highlight the importance of considering tissue-specific biotransformation and the formation of biologically active transformation products to better understand differences in cellular susceptibility to environmental contaminants.

## MATERIALS AND METHODS

### Experimental design

This study combined complementary *in vitro* approaches to investigate 6PPD-Q depletion, distribution, biotransformation, and toxicity using RTgill-W1 and RTBrain cells as model systems. The experimental design included time-resolved exposure experiments, biotransformation profiling, ABCC inhibition experiments, and cytotoxicity assays of 6PPD-Q and selected transformation products and structurally related PPD-quinone analogues following OECD TG 249 assay. Detailed experimental procedures are described in the following sections and in Supplementary Text S1-S5.

### Test systems and chemicals

RTgill-W1 and RTBrain cells (at passages 75-103 and 28-73, respectively), derived from rainbow trout (*Oncorhynchus mykiss*) gill and brain tissues, respectively, were kindly provided by Prof. Bols and colleagues (*75*, *76*). Cells were maintained in Leibovitz L-15 medium supplemented with 5 % fetal bovine serum (FBS) under standard culture conditions described in Supplementary Text S1. Chemicals used in this study are described in Supplementary Table S2. 6PPD-Q was synthesized in-house for Round 1 cytotoxicity experiments (see Supplementary text S5.1) and purchased from HPC Standards® (purity 97.94%) for Round 2 cytotoxicity experiments and biotransformation experiments. Analytical standards of 6PPD-Q biotransformation products and the structurally related PPD-quinone were previously synthesized and characterized by the Peng group at University of Toronto, who employed proton nuclear magnetic resonance (¹H NMR), carbon-13 NMR (¹³C NMR), heteronuclear multiple bond correlation NMR (HMBC-NMR), and liquid chromatography-high resolution mass spectrometry (LC-HRMS) (*30*, *31*, *44*). Additional information on reagents and consumables is provided in Supplementary Text S2.

### Cytotoxicity assessment (OECD TG 249)

Cytotoxicity was assessed according to OECD TG 249 (OECD, 2021)(*77*), using the rainbow trout cell-based multiple-endpoint assay measuring metabolic activity (alamarBlue™), cell membrane integrity (CFDA-AM), and lysosomal integrity (Neutral Red). RTgill-W1 and RTBrain cells were exposed for 24 h to 6PPD-Q, selected BTPs, and related PPD-quinones. Cell viability was expressed relative to the solvent control (0,5% (v/v) DMSO). Detailed information on cell culture conditions, exposure preparation, and endpoint measurements is provided in Supplementary Text S4.

### Time-resolved 6PPD-Q depletion and distribution assay

Time-resolved depletion experiments were conducted in RTgill-W1 and RTBrain cells exposed to 6PPD-Q at a nominal concentration of 1 µg/L in Leibovitz’s L-15 medium supplemented with 5% fetal bovine serum (FBS) following the protocol of Balk et al. (*28*) with minor modifications (*i.e.,* the use of RTBrain cells, lower sampling volume and the use of an automated solid phase extraction protocol). The exposure concentration was selected based on the concentration-response curve for RTBrain cells (Fig. 1B), from which we derived the Non-toxic Concentration (NtC) using the model developed by Stadnicka-Michalak et al. 2018 (*78*). The calculated NtC corresponded to a nominal concentration of 1 µg/L (corresponding to 0.003 µM, value reported in the graphs of the mass balance). Exposure experiments were performed to characterize 6PPD-Q depletion from the medium and its distribution among medium, cellular, and plastic-associated compartments. RTgill-W1 and RTBrain samples were collected at different times over 48 h in three biological replicates, with an additional experiment spanning to 96 h to assess longer-term kinetics. Cell-free controls were included to account for abiotic 6PPD-Q losses, whereas untreated cells (negative controls) were included to monitor cell number over the exposure period, with no 6PPD-Q detected at any sampling time. Apparent *in vitro* clearance was calculated in R over the 0–48 h interval for the three biological replicates from the 48-h experiment and for the additional biological replicate from the extended 96 h experiment, using only measurements collected up to 48 h (Fig. 1G-H). Detailed sampling and extraction procedures are described in Supplementary Text S3.

### Instrumental analysis

Chemical analyses were performed at two different laboratories using laboratory-specific analytical methods and instrumentation. Samples from the cytotoxicity assays conducted in Round 1 (6PPD-Q tested in RTBrain cells) were analyzed at EPFL (Lausanne, Switzerland) using UPLC-MS/MS (Supplementary Text S5.1). All other samples, including those from the biotransformation experiments and cytotoxicity assays conducted in Round 2, were analyzed at Eawag (Dübendorf, Switzerland) using HPLC-HRMS/MS (Supplementary Text S5.2). Sample preparation at Eawag involved automated solid phase extraction (SPE) based on the method reported in Lauper et al. (2022) (*79*). Extracts were analyzed by high-performance liquid chromatography (HPLC; Ultimate, Thermo Fisher Scientific) coupled with high-resolution tandem mass spectrometry (HR-MS/MS; Q-Exactive, Thermo Fisher Scientific).

In the time-resolved depletion experiments, in addition to targeted 6PPD-Q analysis, a literature-based suspect screening analysis was performed to identify potential BTPs (Supplementary Table S1). The identity of selected phase I BTPs was confirmed using analytical standards, whereas phase II TP-O-Gluc was tentatively annotated based on accurate mass. Peak integration of 6PPD-Q and BTPs was performed using Freestyle 1.7 (Thermo Fisher Scientific). 6PPD-Q and phase I BTPs were quantified using compound-specific internal calibration curves, and all samples fell within the respective calibration ranges. TP-O-Gluc was semi-quantified using peak area ratios normalized to the surrogate internal standard, as no synthetic standard was available. Further details about sample preparation and chemical analyses are included as Supplementary Text S3-5.

### Data processing and analysis

From the biotransformation experiments, concentrations of 6PPD-Q measured in the exposure medium, cell-associated fraction, and plastic-associated fraction were converted to absolute mass (µg) using the corresponding extraction volumes and used for mass balance analyses. Furthermore, 6PPD-Q concentrations in the culture medium were plotted against time (h), and first-order elimination rate constants were estimated for each biological replicate. CL_IN VITRO_ values were subsequently calculated after normalization to cell number, following the workflow reported in the OECD 319a test guideline (*80*). CL_IN VITRO_ values reported in the text were calculated in R over the 0–48 h interval for the three biological replicates from the 48-h experiment and for the additional biological replicate from the extended 96 h experiment, using only measurements collected up to 48 h (Fig. 1G-H) (Supplementary Text S3). ChatGPT (GPT-6, OpenAI) was used to assist with writing R scripts for data visualization. Differences in CL_IN VITRO_ between cell lines were assessed using Welch’s t-test. Dose–response curves from the cytotoxicity assays were fitted using nonlinear regression with a four-parameter logistic model in GraphPad Prism (version 5.02; GraphPad Software, San Diego, CA, USA), with Top and Bottom constrained to 100 and 0, respectively. Biological replicates were used for curve fitting, and results are presented as individual replicates, unless otherwise stated. Data processing, mass balance calculations, kinetic analyses, and dose– response analyses were performed using Microsoft Excel, RStudio (version 3.3.0; RStudio version 0.99.902), and GraphPad Prism (version 5.02; GraphPad Software, San Diego, CA, USA).

## Acknowledgment

The authors would like to thank Jessica Bertoli for her valuable support and contribution to the work involving the RTBrain cell line.

## Author contributions

Conceptualization: A.T., K.S.

Methodology: A.T., M.E.F., K.S.

Formal analysis and Investigation: A.T., M.E.F., W.D., T.M., P.N.

Visualization: A.T.

Supervision: K.S.

Writing—original draft: A.T.

Writing—review & editing: A.T., M.E.F., W.D., T.M., P.N., S. G., A. B., E. V., F. B., B.J.D.F., K.S.

Funding acquisition: F.B., B.J.D.F., K.S.

## Use of AI tools

ChatGPT (GPT-6, OpenAI) was used to assist with the translation of manuscript text and to improve its clarity, readability, and grammatical accuracy. The authors reviewed and edited the AI-assisted text and take full responsibility for the final content.

