## Supporting Information for "Tissue-specific biotransformation shapes the toxicity of 6PPD-quinone in fish cell models"

### Supplementary Text.

#### S1. Cell culture conditions.

RTgill-W1 and RTBrain cells were cultured in 150 cm$^2$ cell culture flasks in Leibovitz L-15 medium supplemented with 5 % fetal bovine serum (FBS) at 19 ± 1◦ C in normal atmosphere. When cells reached approximately 90% confluency (~7 and ~10 days for RTgill-W1 and RTBrain, respectively), they were washed twice with Versene and detached using 0.25% trypsin. Following trypsinization, cells were resuspended in L-15/FBS medium, centrifuged for 3 min at 1000 × g at 19 ± 1 °C, and counted using an electronic cell counting system (CASY TCC, Biovendis Products GmbH). Cell suspensions were subsequently diluted in L-15/FBS to obtain the appropriate seeding densities for each experimental setup.

#### S2. Details of Chemicals and Regents.

Leibovitz L-15 medium and trypsin solution were purchased from PAN Biotech, FBS from Eurobio Scientific, and Versene from GibcoTM. Cell culture flasks were obtained from Techno Plastic Product AG, and 24-well culture plates from Greiner Bio-One. Fluorescent dyes used for OECD TG249 assays were purchased from Invitrogen (alamarBlue™ and CFDA-AM) and Sigma-Aldrich (Neutral Red). Details on chemicals tested are reported in Supplementary Table S2.

#### S3. Time-resolved 6PPD-Q depletion and distribution assay

Initial seeding densities were 4.6 × 10$^6$ cells per flask for RTgill-W1 and 2.3 × 10$^6$ cells per flask for RTBrain, using 25-cm$^2$ flask (*29*). RTgill-W1 cells were sampled at 0, 4, 24, and 48 h, whereas RTBrain cells were sampled at 0, 2, 4, 24, and 48 h in three independent biological replicates (n=3), using cells at different passages (Fig. 1D-E). An additional experiment extending the exposure period to 96 h and sampling at 0, 4, 24, 48, 72 and 96 h was conducted in both cell lines (Supplementary Fig. S3). At each sampling point, 1.5 mL of exposure medium was collected, cells were harvested by two successive additions of 750 µL trypsin, and the plastic-associated fraction was recovered by extracting the flask surface twice with 750 µL methanol under gentle agitation for 5 min. Samples were transferred to glass vials and stored at −20 °C until further chemical analysis. Cell-free and negative controls (untreated cells) were subjected to the same sampling procedure as the full treatment. This included also the collection of the fraction corresponding to the cell-associated compartment even if cells were not present. As such, 6PPD-Q detected in this fraction in the absence of cells may reflect residual compound from the exposure medium. Negative controls were also used to count the cell number using the electronic cell counting system CASY TCC (Biovendis Products GmbH) at the beginning and at the end of the exposure period. These measurements were subsequently used to calculate clearance values. Apparent *in vitro* clearance was calculated in R over the 0–48 h interval for the three biological replicates from the 48-h experiment and for the additional biological replicate from the extended 96 h experiment, using only measurements collected up to 48 h (Fig. 1G-H). The difference in apparent clearance between RTBrain and RTgill-W1 was statistically significant (Welch's two-sample t-test, $t = 4.49$, $df = 3.02$, $p = 0.0202$; mean difference = 0.0669 mL $h^{-1}$ $10^6$ $cells^{-1}$, 95% CI: 0.0197–0.1141; $n = 4$ biological replicates per cell line) (Fig. 1I). Chemical analysis of samples from the biotransformation experiments is described in Supplementary Text S5.2.

### S4. Cytotoxicity assay (OECD TG 249)

Cytotoxicity assays were conducted in two independent experimental rounds at different times, following the same experimental approach in accordance with OECD TG 249. Round 1 focused on 6PPD-Q in RTbrain cells (Fig. 1B), whereas Round 2 focused on 6PPD-Q BTPs (Figs. 2B-C, 2E-F) and the other PPD-quinones (Supplementary Figs. S12-13) in both RTbrain and RTgill-W1 cells. In both rounds cell viability was assessed using a multiple-endpoint cytotoxicity assay measuring metabolic activity (alamarBlueTM; AB), cell membrane integrity (5-carboxyfluorescein diacetate acetoxymethyl ester; CFDA-AM) and lysosomal integrity (Neutral Red; NR). Exposures were performed in L-15/ex, which has the same salt and glucose composition as Leibovitz's L-15 medium but does not contain amino acids and vitamins, as previously described by Schirmer et al. (1997) (*81*).

Cell suspensions were diluted in culture medium (L-15 + 5% FBS) and seeded into 24-well plates at densities of 350,000 cells/mL (RTgill-W1) and 150,000 cells/mL (RTBrain) after counting with the electronic cell counting system CASY TCC. Cells were incubated for 48 h at 19 ± 1 °C to allow formation of a confluent monolayer prior to exposure. Serial dilutions of test compounds were prepared from stock solutions in MeOH:DMSO, except for 6PPD-Q, DPPD-Q and DTPD-Q, which were prepared in DMSO only, according to the solvent used for the original stock solutions (Supplementary Table S2). Exposure solutions were prepared by diluting stock solutions in L-15/ex medium while maintaining the final solvent concentration at 0.5% (v/v). Before exposure, cells were washed once with 1 mL L-15/ex to remove residual L-15/FBS. Cells were then exposed to 2.5 mL of the respective dosing solutions.

Immediately after exposure, 500 µL of exposure medium was collected (samples t0), pooled across replicate wells for each concentration, and stored at −20 °C until chemical analysis with HPLC-HRMS (Supplementary Text S5). After 24 h incubation at 19 ± 1 °C in the dark, 500 µL of exposure medium was collected (samples t24) for chemical analysis and similarly pooled, and the remaining exposure solution was removed prior to performing the cell viability assay.

For the multiple-endpoint cytotoxicity assay, cells were washed with 1 mL phosphate-buffered saline (PBS) and incubated for 30 min with 400 µL PBS containing 5% AB (v/v) and CFDA-AM. Fluorescence measurements were performed using a Tecan Infinite M200 plate reader at excitation/emission wavelengths of 530/595 nm for AB and 493/541 nm for CFDA-AM. Following removal of the AB/CFDA-AM solution, cells were incubated with 400 µL PBS containing 1.5% (v/v) NR for 1 h in the dark at 19 ± 1 °C. Cells were then washed with 400 µL fixative solution (0.5% v/v formaldehyde and 1% w/v $CaCl_2$) and incubated with 400 µL extraction solution (1% v/v acetic acid and 50% v/v ethanol) for 10 min in the dark at room temperature under gentle horizontal agitation. NR fluorescence was measured at an excitation/emission wavelength of 530/645 nm. Results were expressed as percentage viability relative to solvent controls based on fluorescence units (FU). Chemical analysis of exposure solutions is described in Supplementary Texts S5.1 and S5.2 for Round 1 and Round 2, respectively.

*S4.1 OECD TG 249 performance*

In absence of specific positive control for RTBrain cells, in Round 1 of OECD TG 249 testing, 3,4-dichloroaniline (3,4-DCA) was preliminary established as such. Therefore, its cytotoxicity was assessed in a preliminary experiment before the main experiments. The resulting values fell within the range specified in OECD TG 249 for RTgill-W1 (*78*) (Supplementary Tables S5 and S6; Supplementary Fig. S16).

In Round 2, assay performance was verified using sodium dodecyl sulfate (SDS) as positive control, recently proposed as less hazardous alternative to 3,4-DCA (*82*). In order to produce additional data for SDS establishment as positive control, in particular for the RTBrain cells, SDS testing was included in all biological replicates for *para*-aryl-OH-6PPD-Q and in at least one biological replicate for all other tested chemicals in both cell lines. SDS concentration–response curves and $EC_{50}$ values in RTgill-W1 were overall consistent with historical laboratory performance and newly established $EC_{50}$ ranges (Supplementary Table S5 and S6 and Supplementary Fig. S16) (*82*). Minor deviations from the new established acceptance ranges were occasionally observed for individual endpoints (CFDA-AM or NR) in some experimental batches (Supplementary Table S5, Fig. S16). However, all three endpoints consistently showed a concentration–dependent decrease in all experiments (Supplementary Fig. S16). Importantly, the most sensitive endpoint (AB) consistently remained within the acceptance range, and at least two of the three endpoints always met the acceptance criteria in every experiment (Supplementary Table S5 and S6). As no established acceptance range was available for SDS in RTBrain cells, the concentration–response curve was adjusted during the experimental campaign, and an RTBrain-specific acceptance range was calculated from all SDS experiments conducted in this study as the mean ± 2.5 SD (Supplementary Table S6), following the approach used to establish the acceptance ranges for SDS in RTgill-W1 (*82*). All positive control results for RTBrain cells, obtained in parallel with the tested compounds, fell within this range (Supplementary Table S5).

All cell viability data generated in this study will be made publicly available through the Fish Cell Acute Toxicity (FACT) database.

*S4.2 ABCC inhibition assay using MK-571.*

To evaluate the potential role of ABCC-mediated efflux during 6PPD-Q exposure, the OECD TG 249 assay with 6PPD-Q was performed in co-exposure with MK-571 at a concentration of 2 µM. 6PPD-Q was tested at nominal concentrations ranging 1-1000 µg/L in three independent biological replicates (Supplementary Fig. S14), with SDS positive control included in two of them (Supplementary Table S5 and Supplementary Fig. S16). This MK-571 concentration was selected based on preliminary concentration–response experiments showing no cytotoxicity in RTgill-W1 cells while increasing intracellular CFDA-AM fluorescence, consistent with inhibition of ABCC-mediated efflux (Supplementary Fig. S15). MK-571 was selected as a broad-spectrum inhibitor of ABCC transporters (*83,84*) considering previous reports showing overall higher expression levels of ABCC transporters in RTgill-W1 compared with RTBrain cells (*29*).

### S5. Chemical analysis

Chemical analyses were performed at two different laboratories using laboratory-specific analytical methods and instrumentation. Samples from the cytotoxicity assays conducted in Round 1 (6PPD-Q tested in RTBrain cells) were analyzed at EPFL (Lausanne, Switzerland) using UPLC-MS/MS (Section S5.1). All other samples, including those from the biotransformation experiments and cytotoxicity assays conducted in Round 2, were analyzed at Eawag (Dübendorf, Switzerland) using HPLC-HRMS/MS (Section S5.2).

*S5.1 UPLC-MS/MS analysis performed at EPFL*

After thawing, a 0.25 mL aliquot of each sample was spiked with an appropriate amount of deuterated internal standard (6PPD-quinone-d5; HPC Standards®), followed by the addition of 0.25 mL of mobile phase (0.1 % formic acid in 95 % water and 5 % acetonitrile). Calibration standards for 6PPD-Q were used for quantification (Supplementary Table S4). UPLC-MSMS analyses were carried out with an ACQUITY UPLC system that was coupled to a Xevo TQ MS mass spectrometer equipped with an electrospray ionization source (ESI) (Waters Corp., Milford, USA). Chromatographic separation was performed with an ACQUITY UPLC HSS T3 column (100 × 2.1 mm, 1.8 µm) at a flow rate of 0.40 mL min-1 and a column temperature of 30 °C. The mobile phase consisted of (A) 0.1 % formic acid in 95 % water and 5 % acetonitrile and (B) 0.1 % formic acid in 95 % acetonitrile and 5 % water. The MS parameters were set as follows; electrospray in positive mode, source temperature: 150°C, desolvation temperature 600°C, cone gas flow (N2): 50 L h-1, desolvation gas flow: 1000 L h-1, collision gas flow: 0.15 mL min-1, ion spray voltage: 1500 V, cone voltage: 30 V. The optimal instrumental parameters were obtained by tuning using direct infusion. Quantitative analyses were performed in MS/MS mode. Identification of 6PPD-Q was based on comparison of two MRM transitions and retention times. The specific parameters are: 5.37 retention time (RT), parent ion 299 (m/z), Dwell time 0.050 (s), Cone voltage 28 (V), Collision voltage 30 (V). Data was acquired and processed with MassLynx V4.2 (Waters Corp., Milford, USA).

*QA/QC (Quality Assurance/Quality Control)*

The surrogate internal standard d5-6PPD-Q was added to all samples, and high linearity was obtained for the calibration curve ($R^2 > 0.99$) (Supplementary Table S4), with a lowest calibration level of 0.1 µg/L. Procedural blanks consisted of repeated injections of mobile phase included in each analytical batch during HPLC-MS/MS analysis to monitor potential background contamination.

*Details regarding the synthesis of 6PPD-Q*

As 6PPD-Q was not commercially available when this study began, it was synthesized in-house in the synthesis chemistry laboratory of EPFL following the protocol of Agua et al. (2021) (*85*). The synthesized compound was then characterized by $^{13}C$ Nuclear Magnetic Resonance (NMR) spectroscopy (Figure S1) and LC-MS/MS, and the results were compared with the compound characterization reported by Tian et al. (*7*).When a commercial 6PPD-Q reference standard became available (HPC Standards®), the in-house 6PPD-Q synthetized compound was further characterized by comparing its high-resolution mass spectrum with that of the commercial standard using an Orbitrap exploris 120 mass spectrometer (Thermo-Fisher®) (Figure S2). The results supported the structural identity of the in-house synthesized compound with the commercial reference standard.

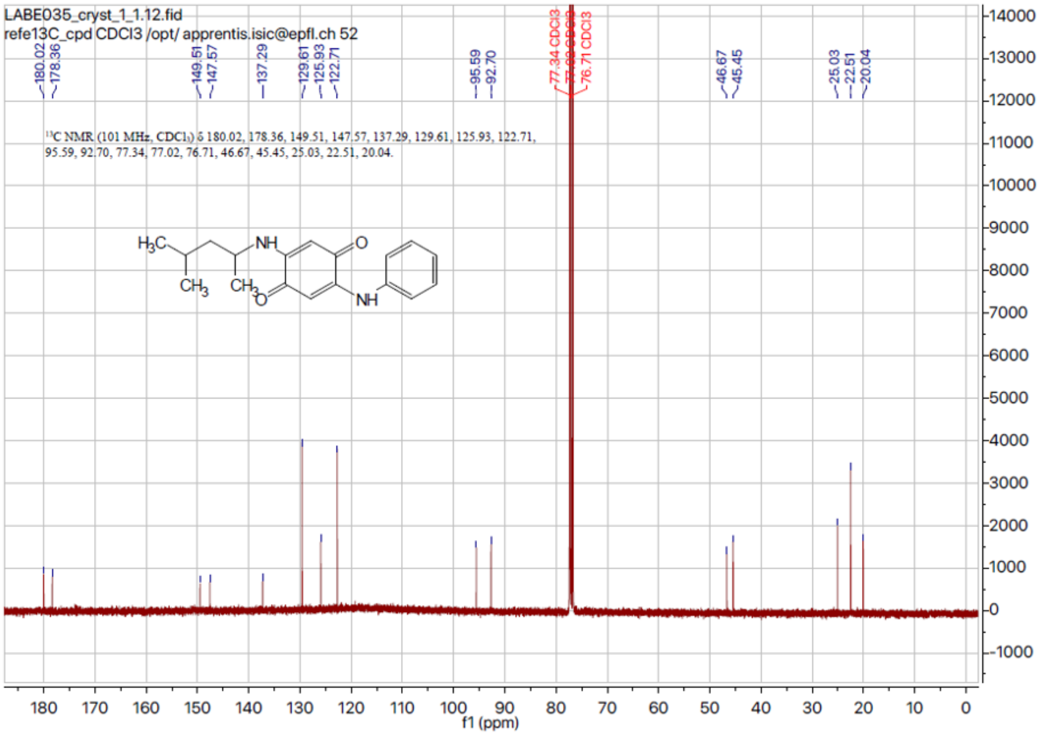

**Figure S1**: $^{13}$C NMR spectrum of the in-house synthesized 6PPD-Q used for Round 1 of cytotoxicity assays.

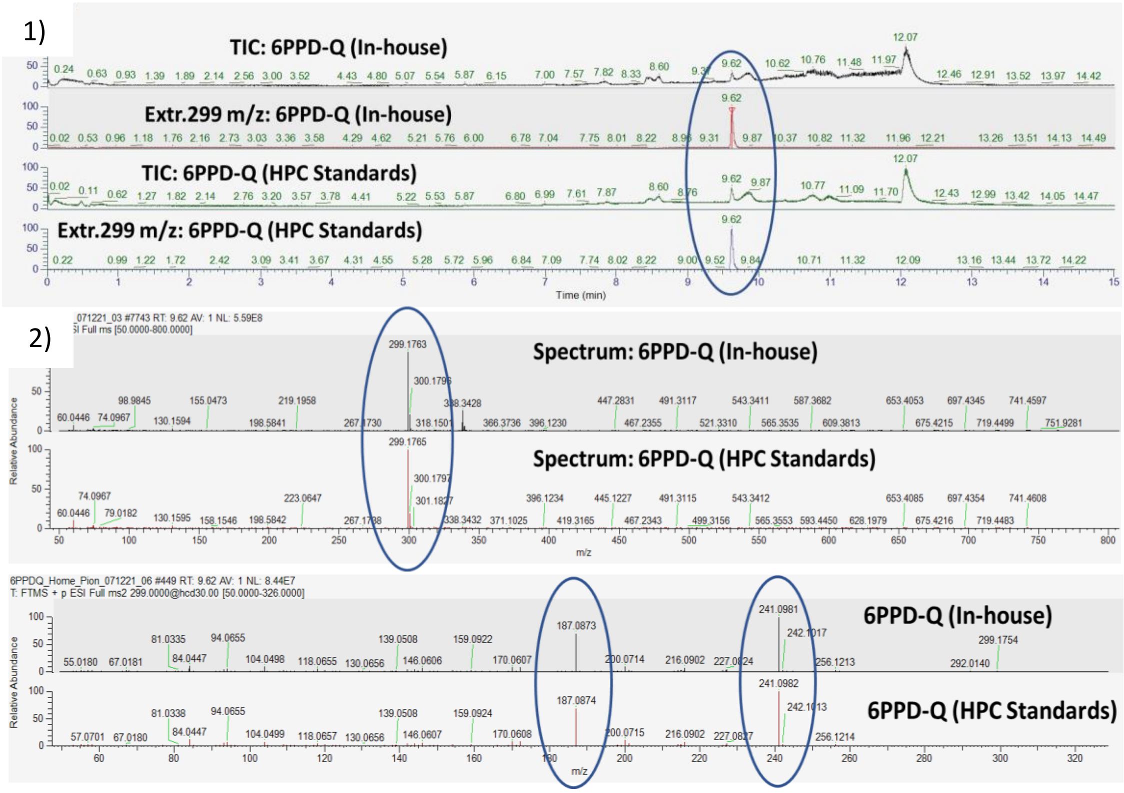

**Figure S2:** 1) Total ion chromatograms (TICs) and extracted ion chromatograms (extr.) for m/z 299 of the in-house synthesized and commercial 6PPD-Q standard (HPC Standards®); 2) high-resolution mass spectra of the precursor ion (MS fragmentation) and products ions (MS2 fragmentation) of both standards.

*S5.2 HPLC-HRMS/MS analysis performed at Eawag*

Samples from the biotransformation experiments and cytotoxicity exposure solutions from Round 2 were extracted using an automated (online) solid phase extraction (SPE) approach (*80*), followed by high-performance liquid chromatography (HPLC; Ultimate, Thermo Fisher Scientific) coupled with high-resolution tandem mass spectrometry (HR-MS/MS; Q-Exactive, Thermo Fisher Scientific).

Briefly, samples from the biotransformation experiments (500 µL) and from the cytotoxicity assay (10 µL, 100 µL or 1000 µL, depending on the nominal concentration) were diluted into an aqueous solution to a total volume of 20 mL in 75.5 x 22.5 mm ND20 Headspace crimp vials containing 0.01 µg $L^{-1}$ of deuterated 6PPD-quinone (6PPD-Q-d5; HPC Standards®). SPE cartridges were filled with Oasis HLB and an ion exchanger mix of strata X-AX and X-CW (Phenomenex) and Isolute ENV+ (Biotage) in a 1:1:1:5 ratio. The 20 mL samples were loaded by dual injection with a flow rate of 2 mL/min using a 2 mM aqueous ammonium acetate ($NH_4C_2H_3O_2$) solution, and the SPE cartridge was eluted with methanol (with 0.1% formic acid) in back flush mode at 40 mL/min prior to chromatography.

HPLC conditions included a flow rate of 0.3 mL/min with the mobile phases A: nanopure water with 0.1 % formic acid and B: methanol also with 0.1% formic acid. The run time per sample was

35 min, with a gradient of 0-20 min 87/13 (A/B), 20-29.5 min 5/95 (A/B), and 29.5-35 min 87/13 (A/B). Chromatographic separation was performed with an Atlantis® T3 (5µm, 3.0 x 150 mm) column. Full scan acquisition included a 100-1000 m/*z* range and a resolution of 140,000 at m/*z* 200, with electrospray ionization in positive mode (ESI+).

6PPD-Q, biotransformation products (BTPs) and the other PPD-quinones were quantified using compound-specific internal calibration curves constructed with synthesized reference standards provided by Hui Peng's group. Details of the standard curves used for quantification are reported in Supplementary Table S4.

Measured concentrations (µg/L) of 6PPD-Q from the biotransformation experiments were converted to absolute amounts (µg) using the corresponding extraction volumes as part of the mass balance approach. Furthermore, a suspect screening workflow combining non-target feature extraction, literature-guided *m/z* searching, and comparison of exposed and control samples was used to identify BTPs (Supplementary Table S1). Putative BTPs were initially selected based on expected *m/z* values reported in the literature and subsequently confirmed by comparison with synthesized reference standards (*ortho*-, *meta*-, and *para*-aryl-OH-6PPD-Q, and C3- and C4-alkyl-OH-6PPD-Q) through retention time matching and chromatographic co-elution (Supplementary Figs. S4-S8, Table S2). Calibration ranges were optimized for each analyte to ensure linear detector response over the concentration range tested, with all samples within the respective calibration ranges. For time-course plots (Supplementary Figs. S9-10), BTP concentrations were reported only for time points at which at least two of the three biological replicates yielded quantifiable concentrations. Values below the lowest calibration level were considered below the quantifiable range and therefore not plotted. In contrast, the transformation product of phase II (TP-O-Gluc) was tentatively annotated based on its accurate mass, as its low abundance precluded MS/MS fragmentation spectra acquisition and no synthetic standard was available for identity confirmation. TP-O-Gluc was not absolutely quantified and was therefore reported as peak area ratios normalized to the surrogate internal standard (Area compound/Area IS).

Measured concentrations of synthesized hydroxylated metabolites (*ortho*-, *meta*-, and *para*-aryl-OH-6PPD-Q, as well as C3- and C4-alkyl-OH-6PPD-Q) and related PPD-quinones (IPPD-Q, CPPD-Q, HPPD-Q, OPPD-Q, DPPD-Q, and DTPD-Q) from the cytotoxicity assays were reported as the geometric mean of t0 and t24 concentrations. Values below the lowest calibration level (0.0039 µg/L in the online vials) were considered below the quantifiable range; for calculation of the geometric mean, these values were assigned a value equal to half of the lowest calibration level according to OECD GD23 (*86*).

*QA/QC (Quality Assurance/Quality Control)*

The surrogate internal standard d5-6PPD-Q was added to all samples to correct for instrumental variability. High linearity was obtained for all calibration curves ($R^2 > 0.97$) (Supplementary Table S4). Instrument performance was monitored throughout the analytical sequence to verify calibration stability and linearity. Procedural blanks consisting of 20 mL Milli-Q water without samples were included in each analytical batch during HPLC-HRMS analysis to monitor potential background contamination. The lowest calibration level in the online vials was 0.0039 µg/L.

**Supplementary Figures.**

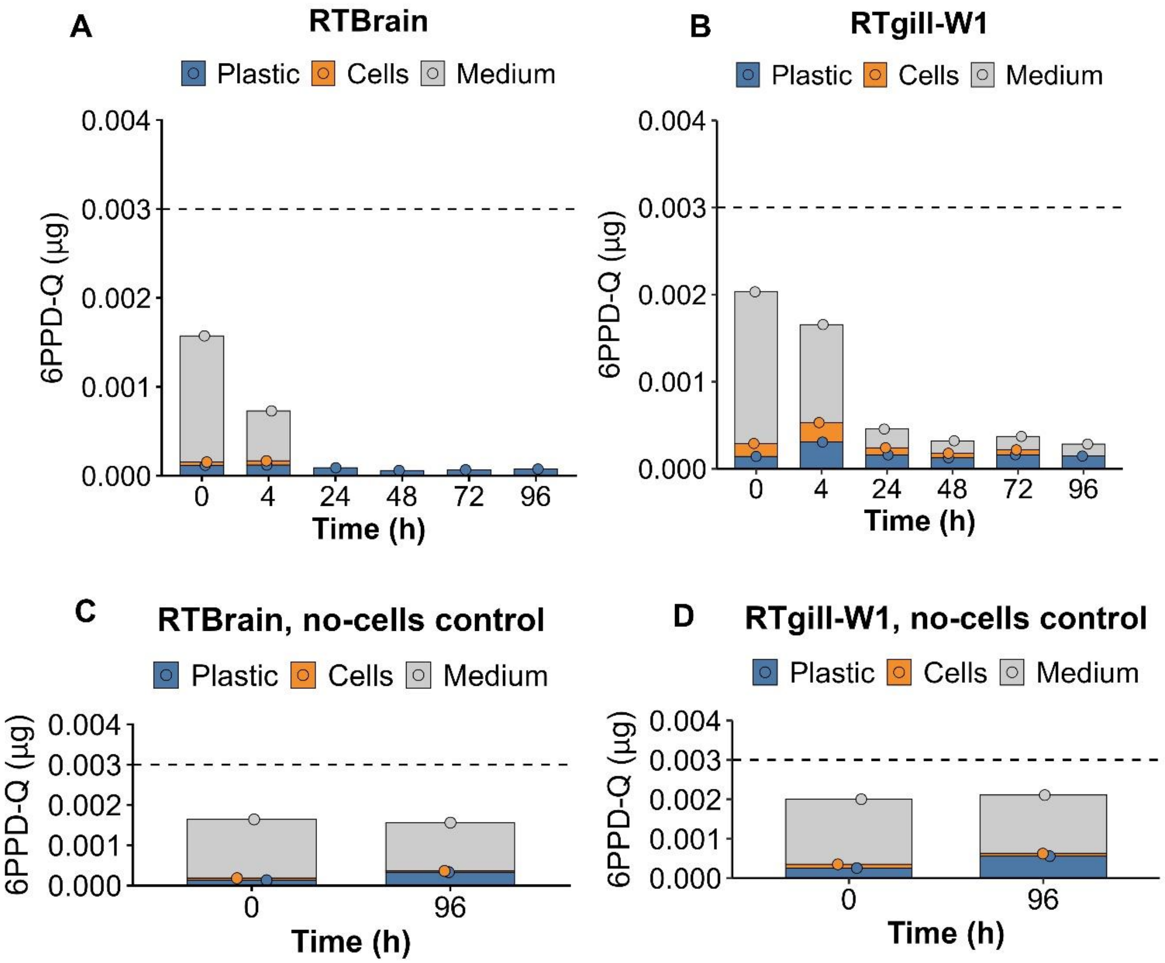

**Figure S3.** (A-B) Time-dependent distribution of 6PPD-Q (μg) across the medium, cellular, and plastic-associated compartments in RTgill-W1 (A) and RTBrain (B) cultures over 96 h from a biological replicate. (C-D) Cell-free control for RTgill-W1 (C) and RTBrain (D) at T0 and 96 h. The same fractionation procedure was applied as for cell-containing cultures to facilitate direct comparison. Consequently, 6PPD-Q detected in this fraction in the absence of cells may reflect residual exposure medium.

In all panels, amounts detected in the medium, cells, and plastic are shown in gray, orange, and blue, respectively. The dashed line indicates the nominal initial mass of 6PPD-Q (0.003 μg), corresponding to the target exposure concentration of 1 μg/L in the 3-mL exposure volume.

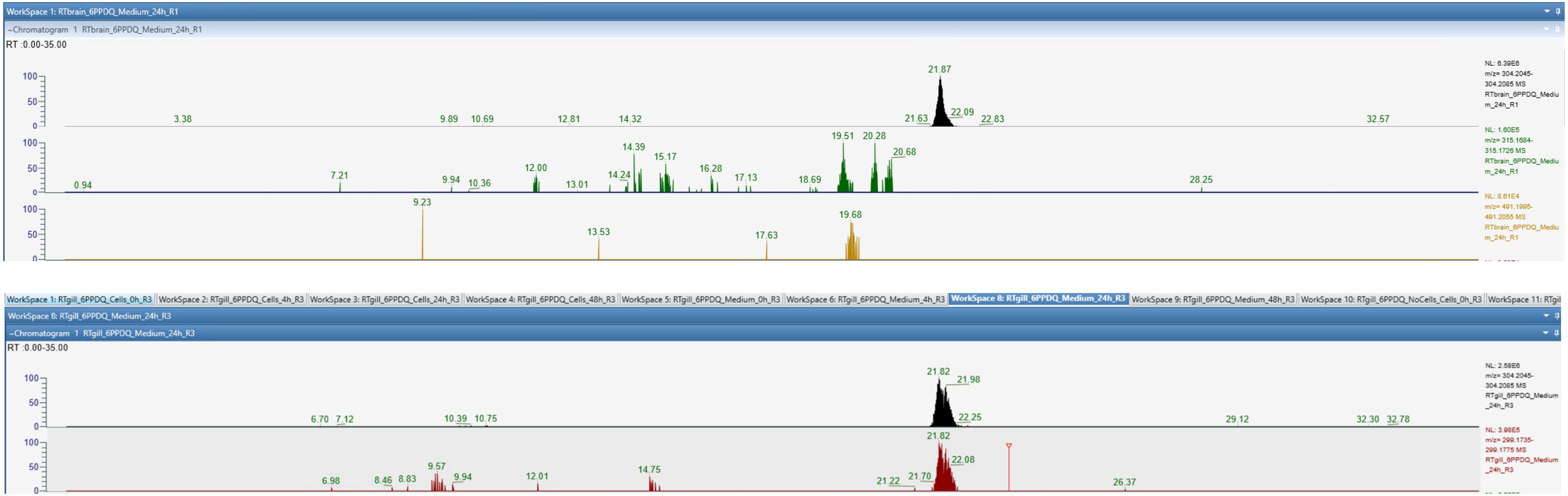

**Figure S4.** Representative extracted ion chromatograms (EICs) of three monohydroxylated compounds (m/z 315.17; green), one glucuronide-conjugated monohydroxylated compound (m/z 491.20; yellow), internal standard (6PPD-Q-d5; black), and 6PPD-Q (red), detected in RTBrain and RTgill-W1 cell extracts. Retention times (RT) are indicated for all detected peaks.

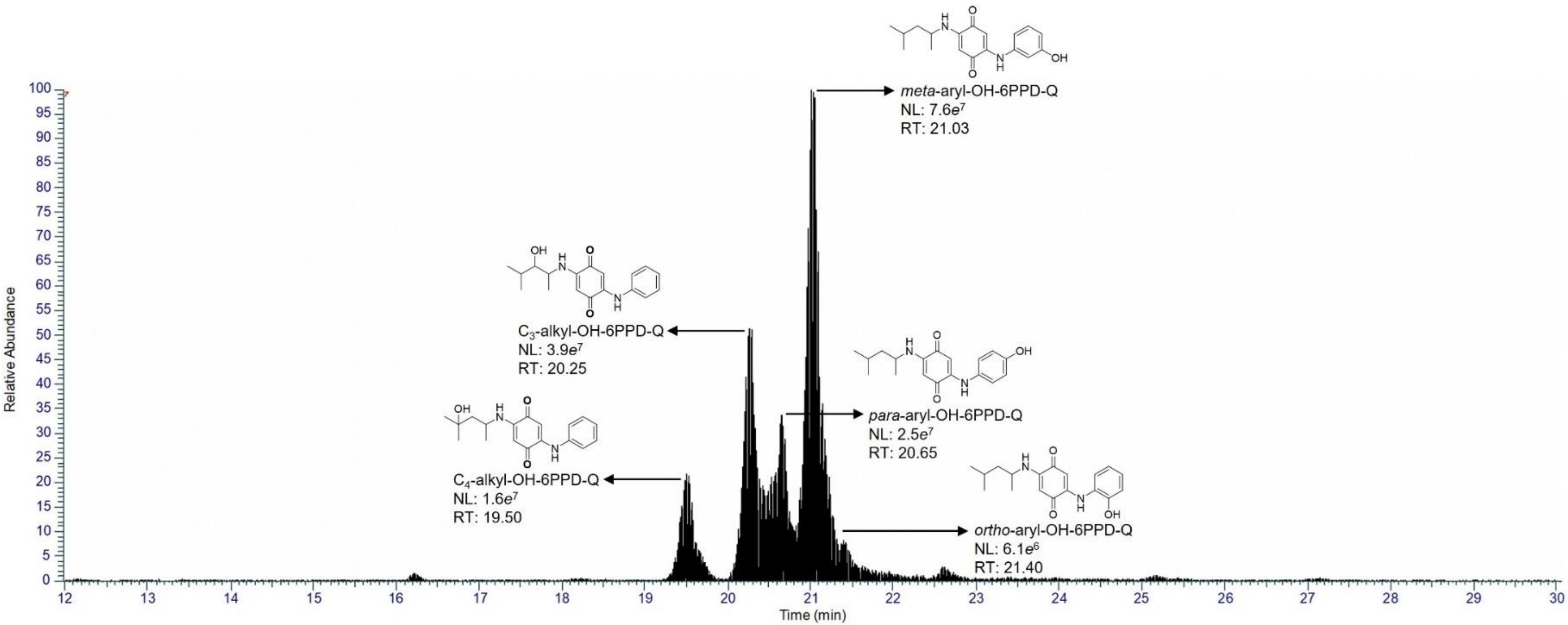

**Fig. S5.** Extracted ion chromatogram of a standard mixture containing five standards of hydroxylated biotransformation products (BTPs) of 6PPD-Q, each injected at a concentration of 0.5 µg $L^{-1}$. The chromatogram shows the chromatographic separation and retention times (RT) of the individual BTP. Corresponding chemical structures are shown adjacent to each chromatographic peak. Peak intensities are presented as relative abundance.

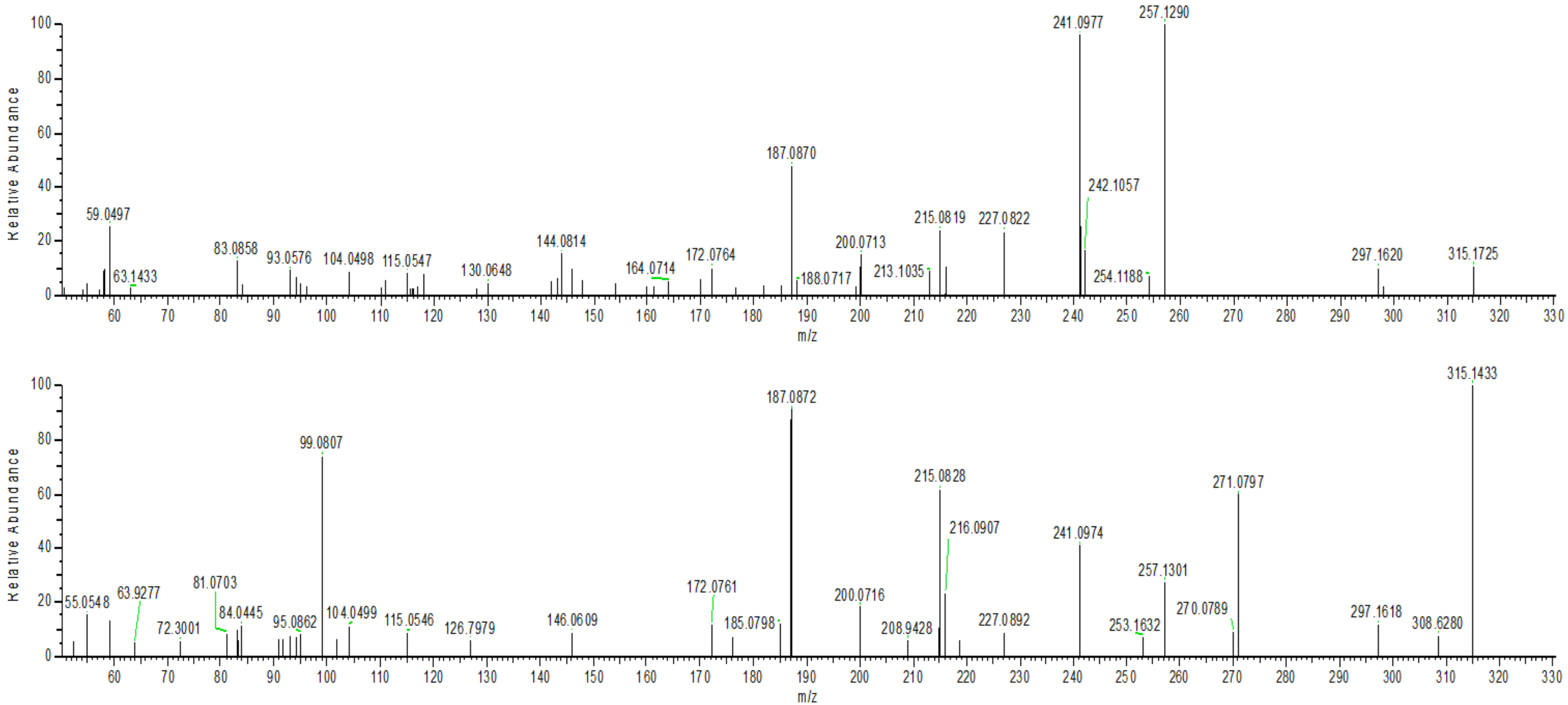

**Fig. S6.** MS/MS spectra of $C_4$-alkyl-OH-6PPD-Q from the synthetic reference standard (top) and a sample from the cell culture medium collected from the cell-based bioaccumulation assessment (bottom). Both spectra exhibit the protonated molecular ion at *m/z* 315.1 and match characteristic fragment ions, including product ions at *m/z* 187.08, 215.08, 241.09 and 257.13, supporting the assignment of $C_4$-alkyl-OH-6PPD-Q in the sample (see Table S1).

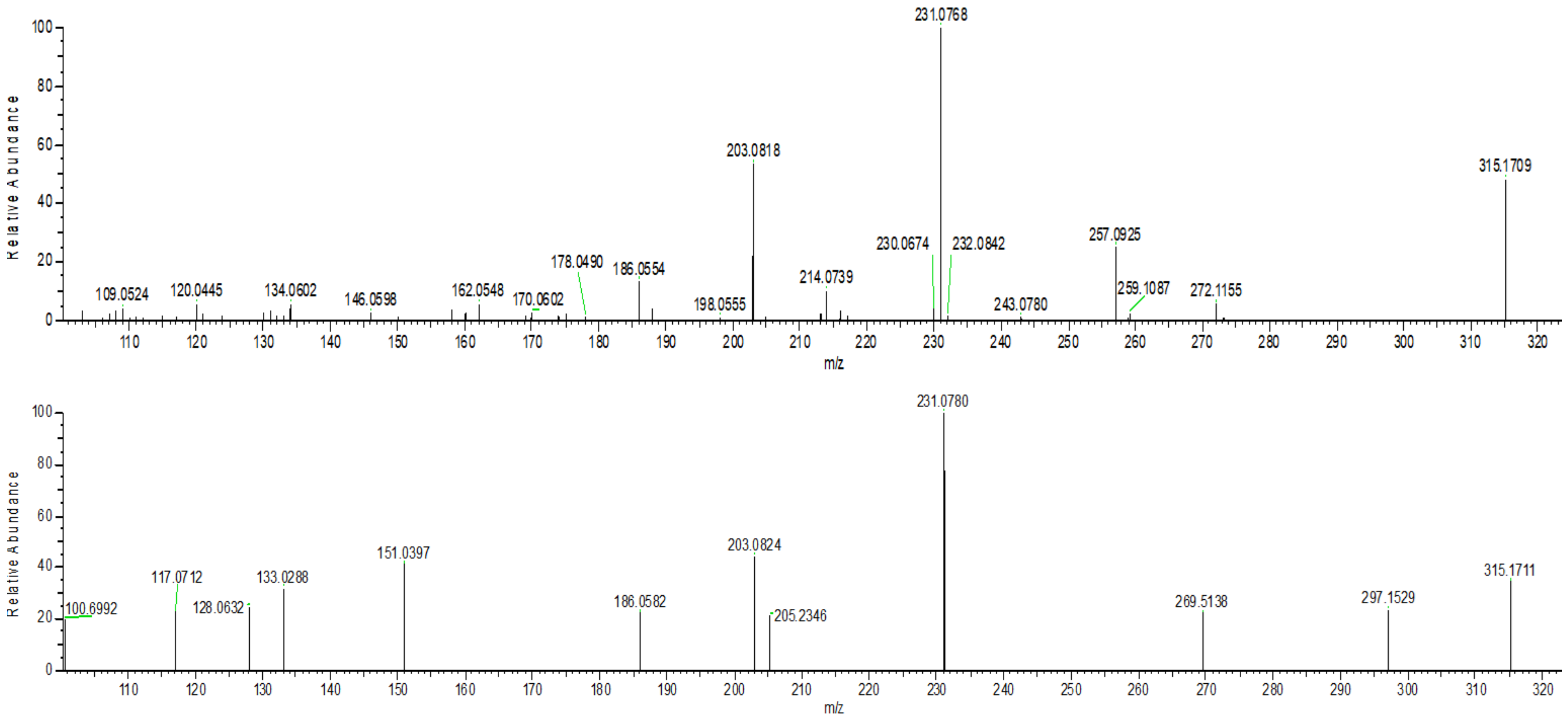

**Fig. S7.** MS/MS spectra of *para*-aryl-OH-6PPD-Q from the synthetic reference standard (top) and a sample from the cell culture medium collected from the cell-based bioaccumulation assessment (bottom). Both spectra exhibit the protonated molecular ion at *m/z* 315.1 and matching characteristic fragment ions, including the dominant product ion at *m/z* 231.08 and the diagnostic fragment at *m/z* 203.08, supporting the assignment of *para*-aryl-OH-6PPD-Q in the sample (see Table S1).

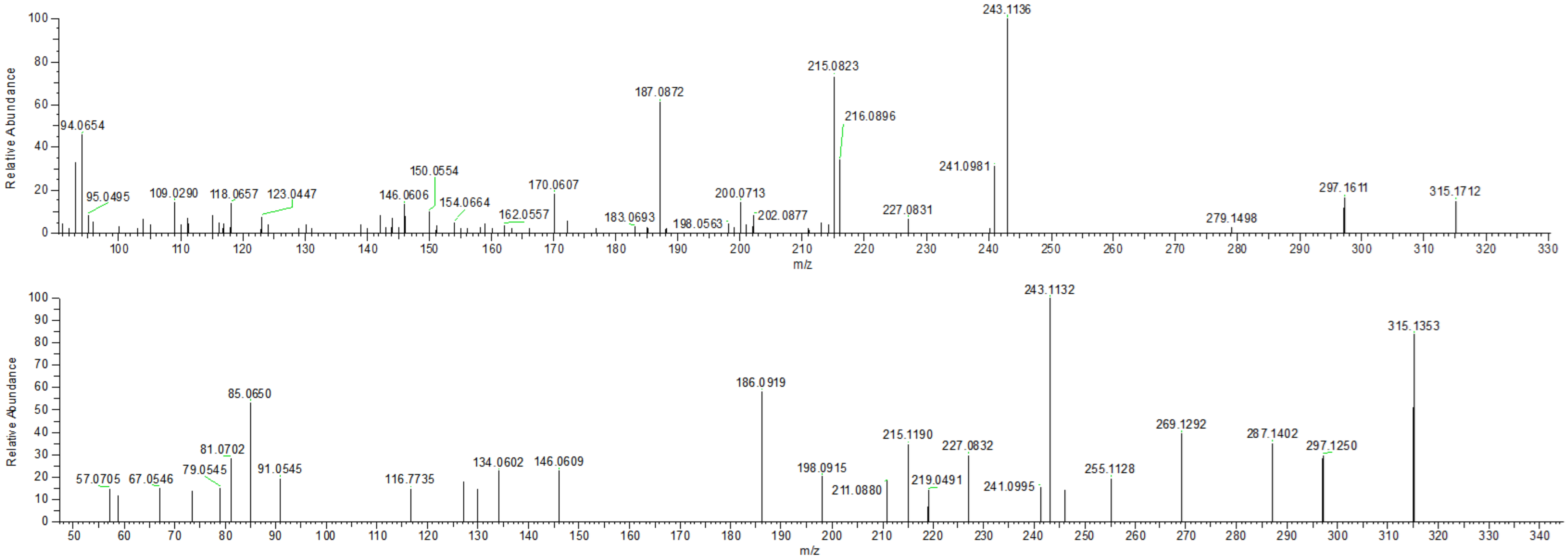

**Fig. S8.** MS/MS spectra of $C_3$-alkyl-OH-6PPD-Q from the synthetic reference standard (top) and a sample from the cell culture medium collected from the cell-based bioaccumulation assessment (bottom). Both spectra exhibit the protonated molecular ion at *m/z* 315.1 and matching characteristic fragment ions, including the dominant product ion at *m/z* 243.1128, supporting the assignment of $C_3$-alkyl-OH-6PPD-Q in the sample (see Table S1).

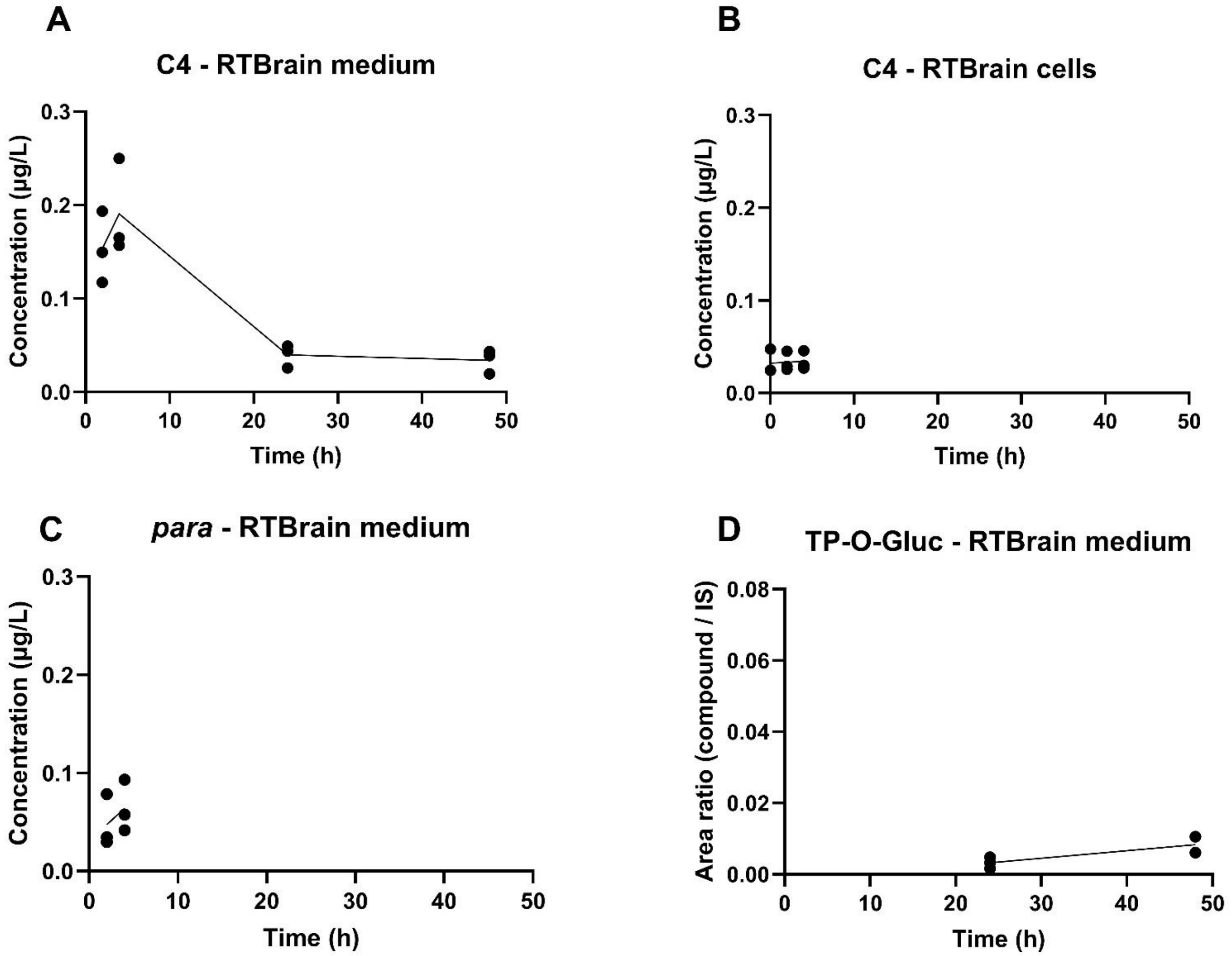

**Figure S9.** Time-dependent formation of 6PPD-Q biotransformation products (BTPs) in RTBrain over a 48-h exposure period (A–D). Points represent independent biological replicates ($n \geq 2$) and lines connect the mean values at each time point. C4-alkyl-6PPD-Q (A, C), and para-aryl-6PPD-Q (B) are expressed as µg/L after quantification using compound-specific calibration curves, whereas TP-O-Gluc (D) is expressed as peak area ratio because no synthetic standard was available.

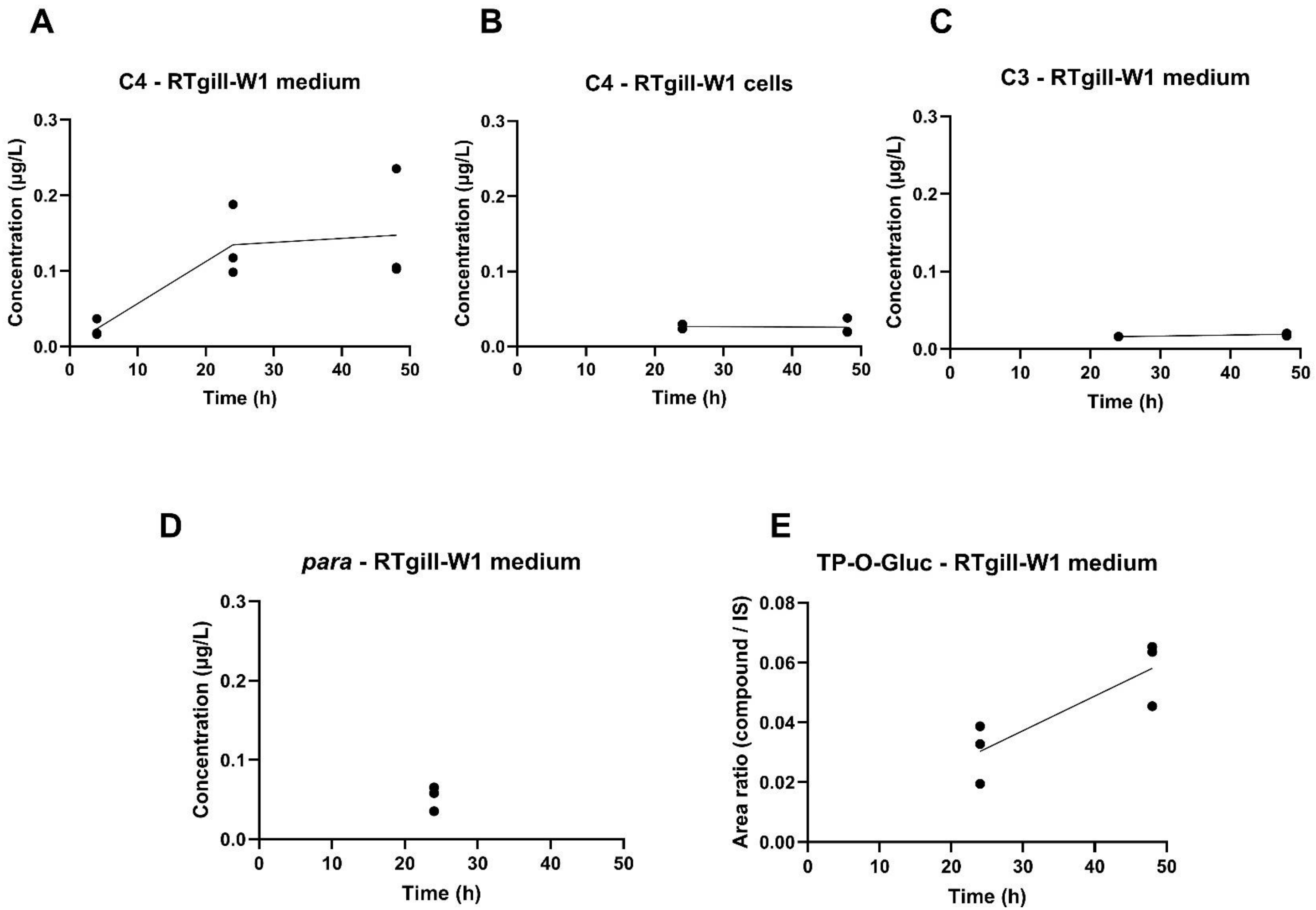

**Figure S10.** Time-dependent formation of 6PPD-Q biotransformation products (BTPs) in RTgill-W1 over a 48-h exposure period (A–E). Points represent independent biological replicates ($n \geq 2$), and lines connect the mean values at each time point. C4-alkyl-6PPD-Q (A, C), para-aryl-6PPD-Q (B), and C3- alkyl-6PPD-Q (D), are expressed as µg/L after quantification using compound-specific calibration curves, whereas TP-O-Gluc (E) is expressed as peak area ratio because no synthetic standard was available.

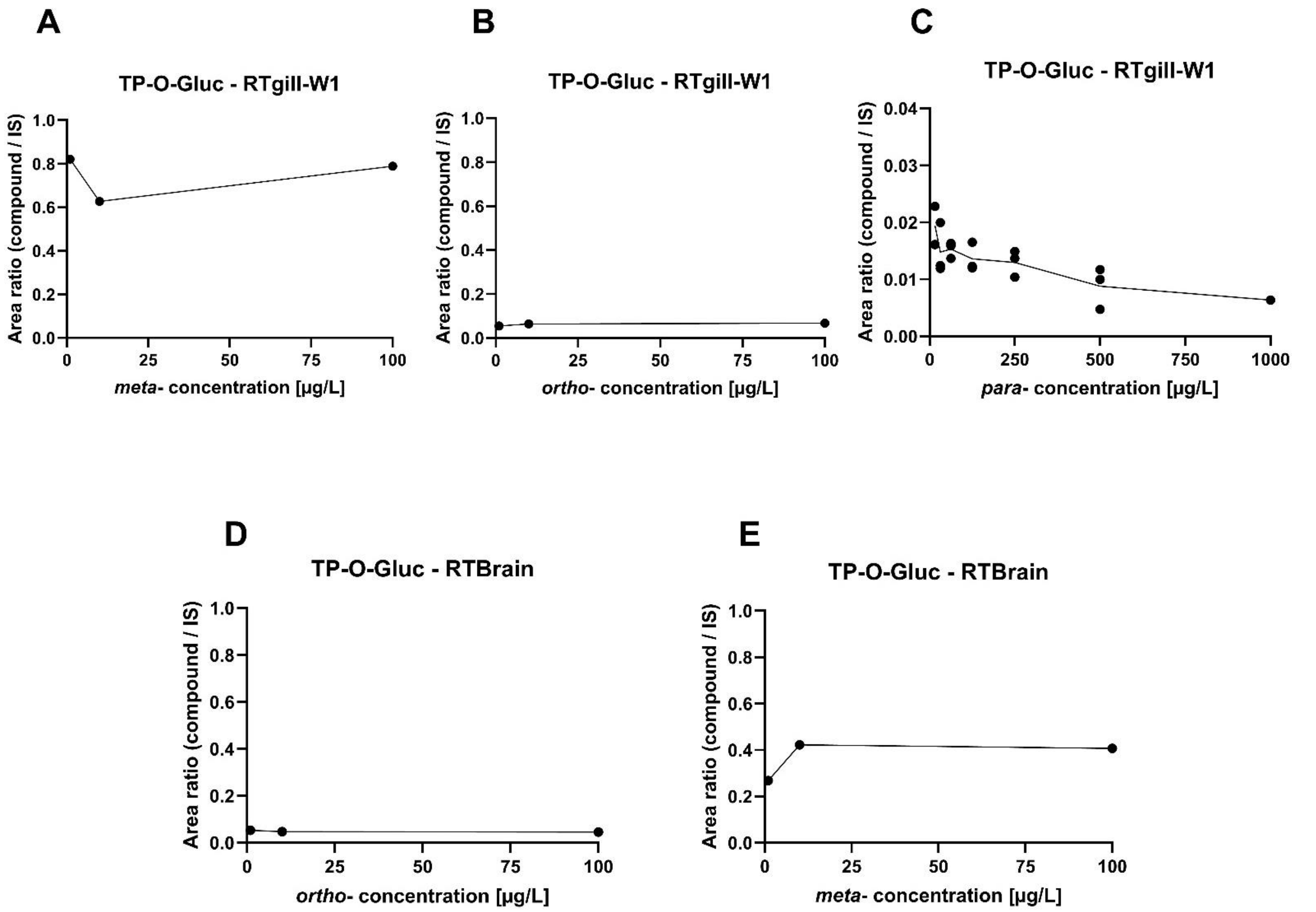

**Figure S11**. Formation of TP-O-Gluc in RTgill-W1 (A–C) and RTBrain (D–E) medium during cytotoxicity assessment under OECD TG 249 following exposure to the synthetic standards *meta*- (A,E), *para*- (C), *ortho*- (B, D) aryl-OH-6PPD-Q. TP-O-Gluc was detected only at 24 h and data are presented as individual biological replicates, with measurement performed in one biological replicate in panels A–B and D–E and in three biological replicates in panel C. No TP-O-Gluc was detected during exposure with C3-, C4-alkyl-OH-6PPD-Q in either cell line or during *para*-aryl-OH-6PPD-Q exposure in RTBrain.

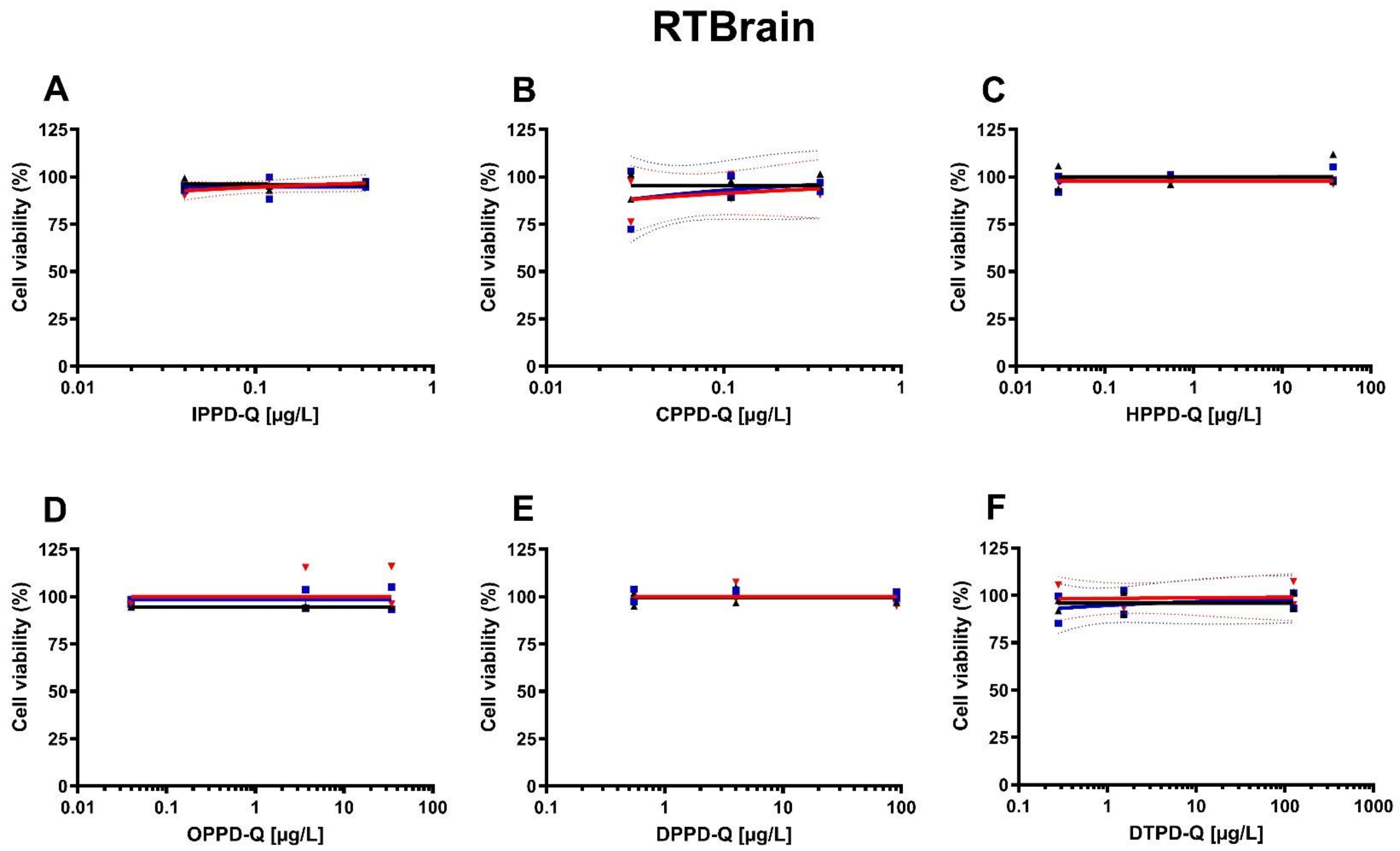

**Figure S12**: Acute cytotoxicity of PPD-Qs in RTBrain cells following OECD TG 249. Dose response curves represent cell metabolic activity (alamarBlue [TM]; blue squares), cell membrane integrity (CFDA-AM; black triangles) and lysosome integrity (Neutral Red; red inverted triangles) after 24 h of exposure. Data are presented as individual replicates (two biological replicates) and expressed as percentage of solvent control (0,5% (v/v) DMSO). Analytical verification of exposure concentrations and SDS positive control were performed in one biological replicate. The resulting exposure concentrations were considered representative of the exposure conditions applied in both biological replicates. No concentration-dependent cytotoxicity was observed within the tested concentration range. Dashed lines indicate the 95% confidence bands around the fitted curves, where calculable.

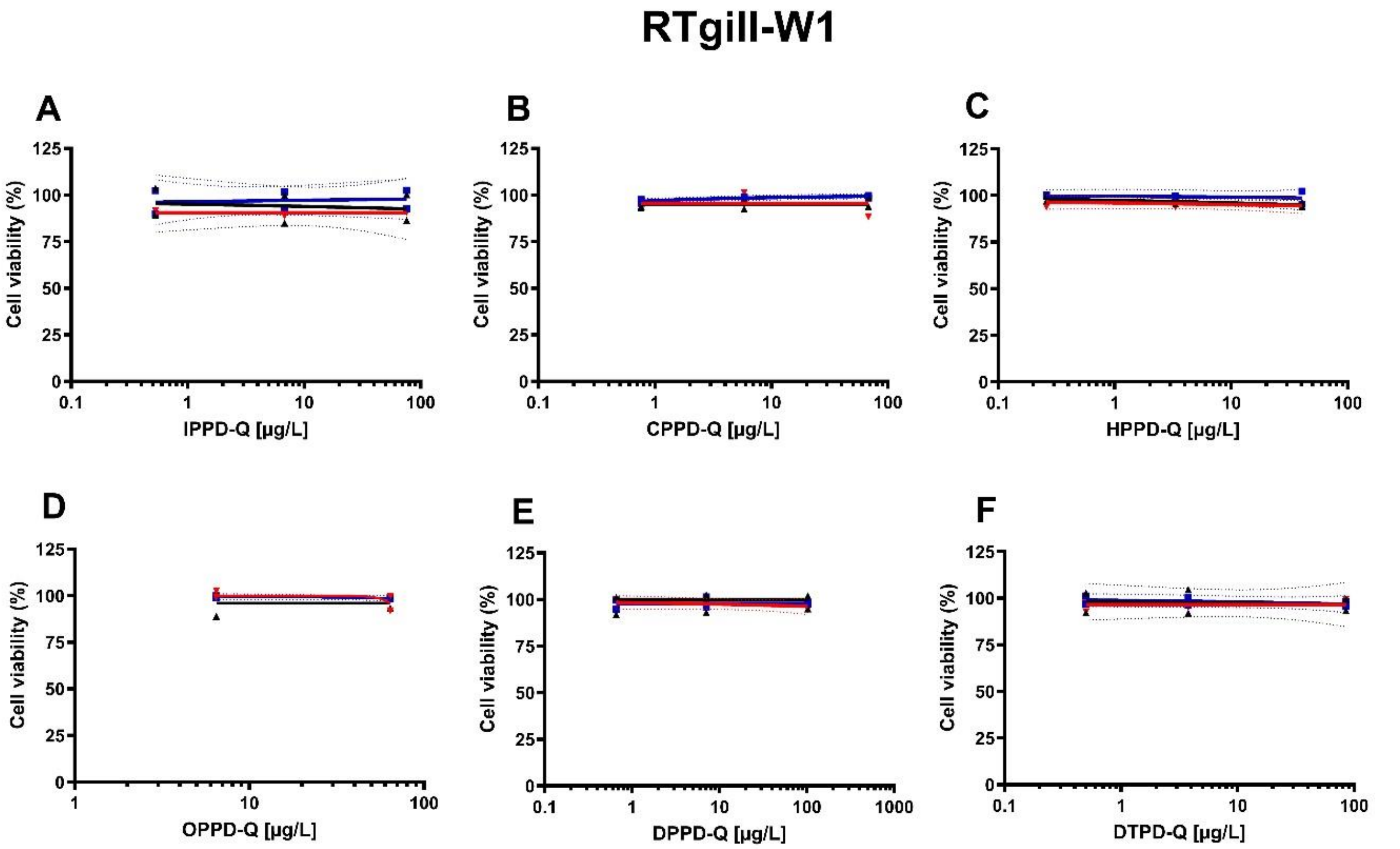

**Figure S13**: Acute cytotoxicity of PPD-Qs in RTgill-W1 cells following OECD TG 249. Dose response curves represent cell metabolic activity (alamarBlue ™; blue squares), cell membrane integrity (CFDA-AM; black triangles) and lysosome integrity (Neutral Red; red inverted triangles) after 24 h of exposure. Data are presented as individual replicates (two biological replicates) and expressed as percentage of solvent control (0,5% (v/v) DMSO). Analytical verification of exposure concentrations and SDS positive control were performed in one biological replicate. The resulting exposure concentrations were considered representative of the exposure conditions applied in both biological replicates. No concentration-dependent cytotoxicity was observed within the tested concentration range. Dashed lines indicate the 95% confidence bands around the fitted curves, where calculable.

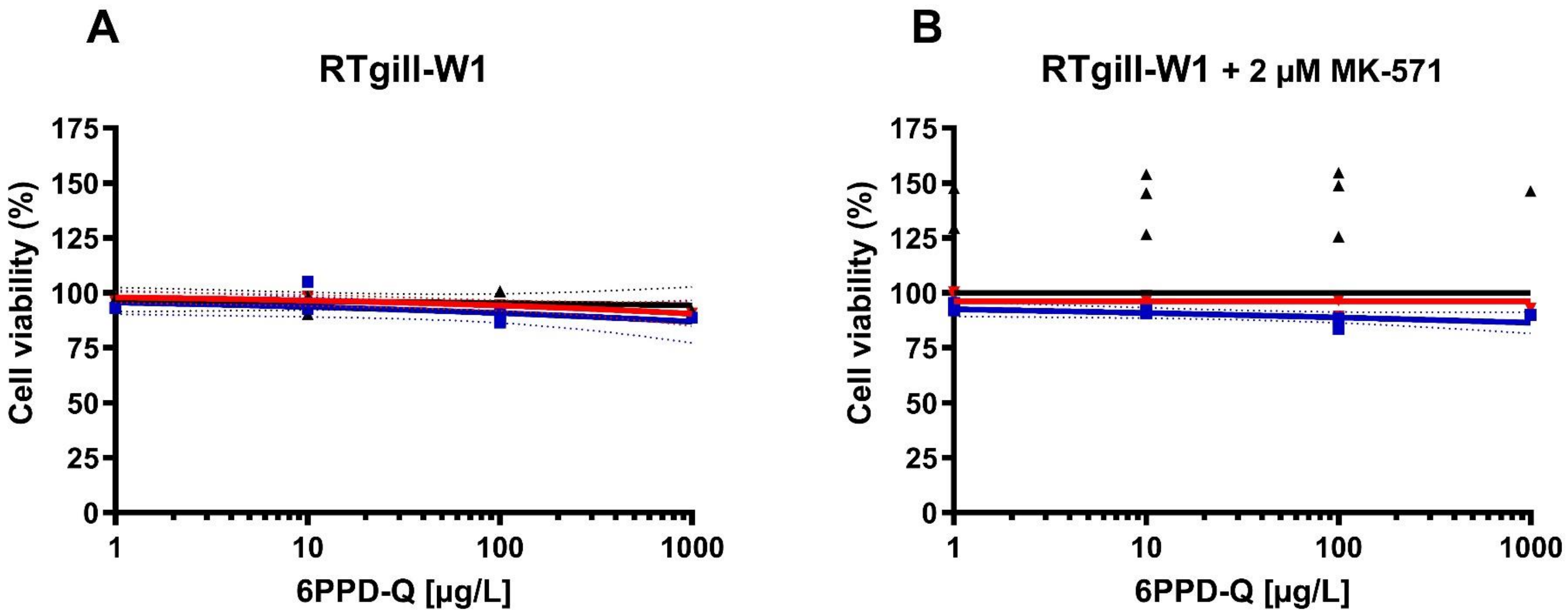

**Figure S14.** Acute cytotoxicity of 6PPD-Q in RTgill-W1 cells following OECD TG 249 without (A) and with MK-571 at 2 µM (B). Dose-response curves represent cell metabolic activity (alamarBlue ™; blue squares), cell membrane integrity (CFDA-AM; black triangles) and lysosomal integrity (Neutral Red; red inverted triangles) after 24 h of exposure. Data are presented individual biological replicates (n = 3) and expressed as percentage of solvent control (0.5% (v/v) DMSO) with SDS positive controls included in two biological replicates. No concentration-dependent cytotoxicity was observed within the tested concentration range. Dashed lines indicate the 95% confidence bands around the fitted curves, where calculable. During dose-response curve fitting, the Top and Bottom values were constrained to 100 and 0, respectively; therefore, the fitted curves do not extend to CFDA-AM values exceeding 100%.

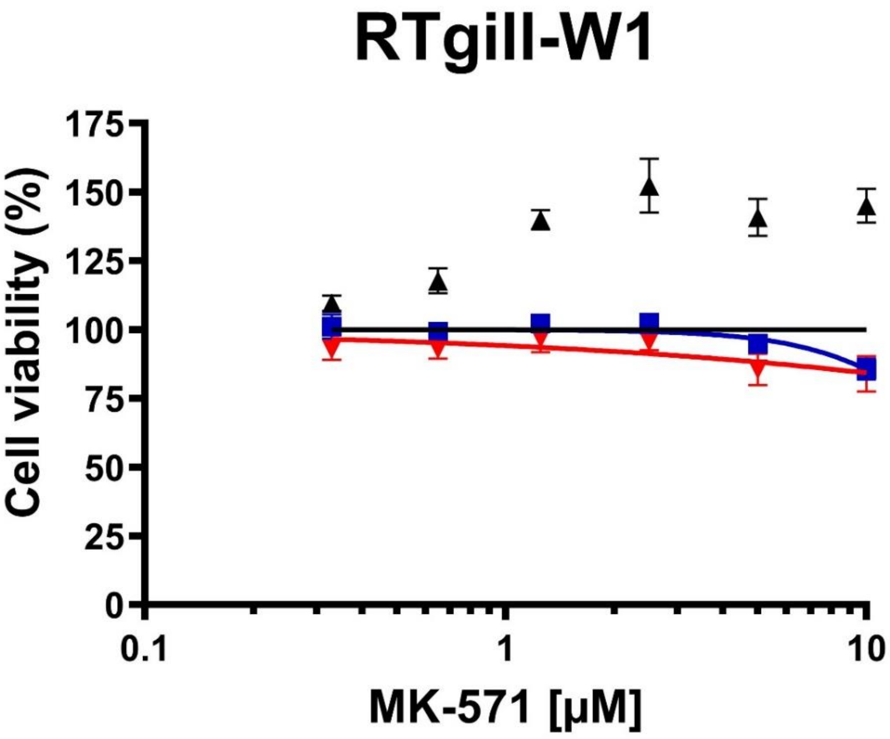

**Figure S15. Dose-response profiles of MK-571 in RTgill-W1 cells as a rationale for concentration selection**. Dose-response curves represent cell metabolic activity (alamarBlue TM; blue squares), cell membrane integrity (CFDA-AM; black triangles) and lysosomal integrity (Neutral Red; red inverted triangles) after 24 h of exposure. The dose-dependent increase in CFDA-AM fluorescence is consistent with accumulation of the hydrolyzed fluorescent dye, carboxyfluorescein, following inhibition of ABCC transporters with the ABCC-targeted inhibitor MK-571. During dose-response curve fitting, the Top and Bottom values were constrained to 100 and 0, respectively; therefore, the fitted curves do not extend to CFDA-AM values exceeding 100%. As shown, RTgill-W1 metabolic activity remained unaffected by MK-571 at concentrations up to 2.5 µM and then decreased. To minimize potential non-specific or cytotoxic effects during subsequent co-exposure, a conservative working concentration of 2 µM was selected for further analyses. Data are expressed as a percentage of the solvent control (0.5% v/v DMSO) and presented as mean ± SD of technical replicates from a single preliminary screening.

**Supplementary Table S1.** Suspect list of 6PPD-Q BTPs compiled from the literature and used for targeted suspect screening of samples from the biotransformation experiment. Reported information includes Compound ID, compound name, chemical formula (if reported), m/z (+), fragment ions (if reported), characteristics, and reference. ***Note:*** *Compound IDs were assigned by the authors. Repeated Compound IDs correspond to the same compound found in different publications.*

| ID | Compound name | Chemical formula (if reported) | m/z (+) | Fragment ions (if reported) | Characteristics | Reference |
|---|---|---|---|---|---|---|
| 0 | 6PPD-Q (CAS 2754428-18-5) | C18H22N2O2 | 299.1750 | 256.121; 241.097; 215.082; 200.071; 187.087; 100.112 | parent | Ankley et al., 2025 |
| 0 | 6PPD-Q-d5 (2750119-14-1) | - | 304.2070 | 261.152; 248.144; 246.129; 220.113; 205.102; 192.118 | parent_d5 | |
| 1 | TP-OH1 or 6PPD-Q-4-OH | - | 315.1700 | 257.129; 241.097; 215.082; 200.071; 187.087; 164.071; 99.081 | monohydroxylated TP on carbon 4 position of the alkyl side chain | |
| 1 | TP-OH2 or p-hydroxy 6PPD-Q (40605) | - | 315.1700 | 272.116; 257.092; 231.077; 203.082 | monohydroxylated TP on the para position of the phenyl ring | |
| 2 | TP-2-OH | - | 331.1650 | 313.154; 273.123; 257.092; 243.077; 231.077; 216.066; 203.082 | dihydroxylated TP | |
| 5 | TP-O-Gluc1 | - | 491.2020 | 315.170; 241.098; 215.082; 187.087 | glucuronide-conjugate monohydroxilated | |
| 5 | TP-O-Gluc2 | - | 491.2020 | 315.170; 231.076; 203.082 | glucuronide-conjugate monohydroxilated | |
| 6 | TP-2-O-Gluc-1 | - | 507.1980 | 331.165; 257.092; 231.077; 203.082 | glucuronide-conjugate dihydroxilated | |
| 6 | TP-2-O-Gluc-2 | - | 507.1980 | 331.165; 231.077; 203.082 | glucuronide-conjugate dihydroxilated | |
| 6 | TP-2-O-Gluc-3 | - | 507.1980 | 331.165; 241.098; 231.077; 215.082; 203.081 | glucuronide-conjugate dihydroxilated | |
| 6 | TP-2-O-Gluc-4 | - | 507.1980 | 331.166; 247.072 | glucuronide-conjugate dihydroxilated | |
| 7 | TP-2-O-Sulf | - | 411.1220 | 411.122; 337.049; 311.034; 257.092; 231.077; 203.082 | sulfonate-conjugated dihydroxilated | |
| 1 | OH-6PPD-Q | C18H23N2O3 | 315.1690 | 94.0651; 99.0809; 187.0862;215.0811; 241.0971; 257.1283 | monohydroxylated TP | Nair et al., 2025 (*Synthesis and Toxicity Evaluation of p-Phenylenediamine Quinones*) |
| 1 | OH-6PPD-Q | C18H23N2O3 | 315.1690 | 110.0601; 203.0811; 231.0761 | monohydroxylated TP | |

| | | | | | | |
|---|---|---|---|---|---|---|
| 0 | 6PPD-Q | - | 299.1760 | - | parent | Nair et al., 2025 (*Structure and Toxicity Characterization of Alkyl Hydroxylated Metabolites of 6PPD-Q*) |
| 1 | alkyl-OH-6PPD-Q (C4) - standard | - | 315.1709 | 241.0969; 257.1284 | monohydroxylated TP | |
| 1 | alkyl-OH-6PPD-Q (C3) - standard | - | 315.1709 | 243.1128 | monohydroxylated TP | |
| 1 | alkyl-OH-6PPD-Q (C4) from fish tissue extract | - | 315.1709 | 241.0969; 257.1284 | monohydroxylated TP | |
| 1 | alkyl-OH-6PPD-Q (C3) from fish tissue extract | - | 315.1709 | 243.1128 | monohydroxylated TP | |
| 1 | alkyl-OH-6PPD-Q (C4) from from liver S9 and microsome | - | 315.1709 | - | monohydroxylated TP | |
| 1 | aryl-OH-6PPD-Q from from liver S9 and microsome | - | 315.1709 | - | monohydroxylated TP | |
| 1 | *para*-aryl-OH-6PPDQ | - | - | - | monohydroxylated TP | Nair et al., 2025 (*Regioselective In Vitro Toxicity of Aromatic Hydroxylated Metabolites of 6PPD-Quinone*) - Chemrxiv |
| 1 | *ortho*-aryl-OH-6PPDQ | . | . | . | monohydroxylated TP | |
| 1 | *meta*-aryl-OH-6PPDQ | . | . | . | monohydroxylated TP | |
| 8 | Bio transform 3.0 (1) | C28H37N5O8S | 603.2362 | - | GSH-conjugation of alpha, beta-unsaturated carbonyl | GSH conjugates identified with Biotransform 3.0 |
| 8 | Bio transform 3.0 (2) | C28H37N5O8S | 603.2362 | - | GSH-conjugation of alpha, beta-unsaturated carbonyl | |
| 9 | Bio transform 3.0 (3) | C28H39N5O8S | 605.2519 | - | GSH-conjugation of quinone | |
| 9 | Bio transform 3.0 (4) | C28H39N5O8S | 605.2519 | - | GSH-conjugation of quinone | |
| 10 | 6QTP297 | C18H20N2O | 297.1598 | 215.0813; 241.0969; 297.1596 | Dehydrogenated 6PPD-Q | Wang et al., 2025 |
| 11 | 6QTP313 | C18H20N2O3 | 313.1547 | 83.0861; 231.0759; 257.0915; 313.1552 | Dehydrogenated monohydroxy 6PPD-Q | |
| 1 | 6QTP315L | C18H22N2O3 | 315.1703 | 83.0861; 187.0863; 241.0966; 315.1688 | Monohydroxy 6PPD-Q | |

| | | | | | |
|---|---|---|---|---|---|
| 1 | 6QTP315M | C18H22N2O3 | 315.1703 | 85.1017; 203.0808; 231.0757; 273.1244; 315.1714 | Monohydroxy 6PPD-Q |
| 1 | 6QTP315R | C18H22N2O3 | 315.1703 | - | Monohydroxy 6PPD-Q |
| 12 | 6QTP329AL | C18H20N2O4 | 329.1496 | 112.0757; 187.0862; 200.0704; 241.0966; 283.1431; 311.1381; 329.1488 | Monohydroxy 6PPD-Q aldehyde |
| 12 | 6QTP329AR | C18H20N2O4 | 329.1496 | 112.0757; 187.0864; 200.0702; 241.0966; 283.1433; 311.1381; 329.1488 | Monohydroxy 6PPD-Q aldehyde |
| 2 | 6QTP331L | C18H22N2O4 | 331.1652 | 59.0494; 83.0860; 108.0450; 188.0705; 203.0813; 231.0760 | Dihydroxy 6PPD-Q |
| 2 | 6QTP331R | C18H22N2O4 | 331.1652 | 75.0447; 187.0863; 215.0811; 241.0968<br>257.1278; 331.1645 | Dihydroxy 6PPD-Q |
| 13 | 6QTP349 | C18H24N2O5 | 349.1758 | 71.0498; 100.0759; 219.1157; 261.1266; 331.1645; 349.1738 | Trihydroxy 6PPD hydroquinone |
| 14 | 6QTP477L | C24H32N2O | 477.2231 | 99.0807; 215.0811; 315.1696;477.2211 | 6PPD-Q O-glucoside |
| 14 | 6QTP477R | C24H32N2O | 477.2231 | 210.1852; 231.0716; 315.1698; 436.1194;477.2231 | 6PPD-Q O-glucoside |
| 15 | 6QTP493 | C24H32N2O9 | 493.2181 | 231.0760; 257.0916; 419.1442; 493.2174 | Monohydroxy 6PPD-Q O-glucoside |
| 16 | 6QTP395L | C18H23N2O6P | 395.1366 | 83.0858; 215.0810; 241.0966; 297.1592;395.1370 | 6PPD-Q phosphoryl |
| 16 | 6QTP395R | C18H23N2O6P | 395.1366 | - | 6PPD-Q phosphoryl |
| 17 | 6QTP411 | C18H23N2O7P | 411.1316 | 81.0703; 215.0814; 241.0975; 295.1430; 393.1202; 411.1292 | Monohydroxy 6PPD-Q phosphoryl |
| 18 | 6QTP317 | C18H24N2O3 | 317.186 | 70.0658; 86.0969; 129.1023; 216.0862;317.1876 | Monohydroxy 6PPD hydroquinone |
| 19 | 6QTP361L | C19H24N2O | 361.1779 | - | Methoxy dihydroxy 6PPD-Q |
| 19 | 6QTP361M | C19H24N2O | 361.1779 | - | Methoxy dihydroxy 6PPD-Q |
| 19 | 6QTP361R | C19H24N2O | 361.1779 | 218.0445; 260.0552; 361.1753 | Methoxy dihydroxy 6PPD-Q |
| 20 | 6QTP218 | C12H11NO | 218.0812 | - | 2-hydroxyl-5-phenylamino-1,4-quinone |

| | | | | | | |
|---|---|---|---|---|---|---|
| 21 | 6QTP329L | C19H24N2O3 | 329.186 | 202.0735; 257.0930; 329.1875 | Methoxy 6PPD-Q | |
| 21 | 6QTP329R | C19H24N2O3 | 329.186 | - | Methoxy 6PPD-Q | |
| 22 | 6QTP347 | C18H22N2O5 | 347.1601 | - | Trihydroxy 6PPD-Q | |
| 23 | 6QTP345L | C19H24N2O | 345.1809 | - | Methoxy monohydroxy 6PPD-Q | |
| 23 | 6QTP345R | C19H24N2O | 345.1809 | 85.0653; 142.0863; 261.1264;345.1837 | Methoxy 6PPD hydroquinone aldehyde | |
| 0 | 6PPD-Q | - | 299.1700 | | parent | Montgomery et al., 2023 |
| 1 | OH-6PPD-Q | - | 315.1700 | 231.08; 315.17 | monohydroxylated TP | |
| 5 | 6PPD-Q-*O*-Gluc | - | 491.2000 | 231.08; 315.17 | glucuronide conjugation | |
| 1 | OH-6PPD-Q | C18H22N2O3 | 315.1703 | 83.0855; 241.0973; 257.1286; 315.1711 | Mono hydroxy-<br>6PPD-Q | Qi et al., 2024 |
| 5 | 6PPD-Q-Ogluc | C24H30N2O9 | 491.2023 | 315.1705; 491.2030 | 6PPD-Q-*O*-glucuronide | |
| 0 | 6PPDQ | C18H25N2 (+) | 299.1760 | - | parent | Grasse et al., 2023 |
| 5 | 6-PPDQ+O+glucuronide | C24H31N2O9 (+) /<br>C24H29N2O9 | 491.2030 | - | hydroxylated + glucuronidated metabolite | |
| 25 | 6-PPDQ+O+SO3 | C18H23N2O6S (+) /<br>C18H21N2O6S | 395.1263 | - | hydroxylated + sulfated metabolite | |
| 6 | 6-PPDQ+2 O+glucuronide | C24H31N2O10 (+) /<br>C24H29N2O10 (-) | 507.1978 | - | dihydroxylated + glucuronidated metabolite | |
| 7 | 6-PPDQ+2O+SO3 | C18H23N2O7S (+) /<br>C18H21N2O7S (-) | 411.1220 | - | dihydroxylated + sulfated metabolite | |
| 26 | 6-PPDQ+O+GSH | C28H38N5O9S (+) /<br>C28H36N5O9S (-) | 620.2386 | - | hydroxylated glutathione conjugate | |
| 27 | 6-<br>PPDHQ+acylglucuronide+NAc<br>Cys | C30H40N3O13S (+) /<br>C30H38N3O13S (-) | 682.2278 | - | complex Phase II conjugate | |
| 28 | 6-PPDQ+2O+CH3+Cys | C22H29N3O6S (+) /<br>C22H28N3O6S (-) | 464.1835 | - | oxidized + methylated + cysteine conjugate | |

| | | | | | | |
|---|---|---|---|---|---|---|
| 29 | 6-PPDQ+O+NAcCys | C23H29N3O6S | 476.1846 | - | hydroxylated + N-acetylcysteine | |
| 1 | 6-PPDQ+O | C18H23N2O3 | 315.1702 | - | monohydroxylated primary metabolite | |
| 1 | monohydroxylated metabolite | C18H23N2O3 | 315.1709 | - | mono-hydroxilated | Hiki & Yamamoto 2022 |
| 0 | 6PPD-Q | - | 299.1754 | 215.0816; 187.0868; 241.0972; 256.1206; 200.0709; 100.1126 | parent | Ankley et al., 2024 |
| 0 | 6PPD-Q_d5 | - | 304.2068 | 220.1131; 192.1180; 246.1286; 261.1521; 205.1022; 248.1443 | IS | |
| 1 | TP-OH1 | - | 315.1703 | 257.1288; 241.0975; 215.0819; 200.0711; 187.0870; 164.071; 99.0811 | mono-hydroxilated compound | |
| 1 | TP-OH2 | - | 315.1703 | 272.1157; 231.0765; 203.0818 | mono-hydroxilated compound | |
| 1 | TP-1 | - | 315.1703 | 243.1131; 216.0898; 200.0710; 150.0552; 94.0658 | mono-hydroxilated compound | |
| 2 | TP-2OH1 | - | 331.1652 | 164.071; 99.0810; 231.0767; 203.0818 | dihydroxylated transformation products | |
| 2 | TP-2OH2 | - | 331.1652 | 225.43; 186.9231; 257.1287; 241.0972; 215.0819 | dihydroxylated transformation products | |
| 2 | TP-2OH3 | - | 331.1652 | 231.0767; 203.08; 216.0658; 150.0552 | dihydroxylated transformation products | |
| 5 | 6PPD-Q glucuronidation | - | 491.2024 | 315.17055; 231.07667 | glucuronidation of 6PPD-Q | |
| D5 | TP-d5-OH1 | - | 320.2018 | 99.0811; 192.1183; 220.1132; 246.1293; 262.1604 | mono-hydroxilated compound | |
| D5 | TP-d5-OH2 | - | 319.1960 | 207.1071; 235.1020; 261.1176 | mono-hydroxilated compound | |
| D5 | TP-d5-OH3 | - | 320.2021 | 208.1133; 236.1082; 262.1238 | mono-hydroxilated compound | |

**Supplementary Table S2.** Analytical standards used in this study, including their chemical structures, molecular formulas, stock solution concentrations, retention times (RT), and mass-to-charge ratios (m/z) in positive ionization mode (+).

| Standard Name | Structure | Molecular formula | Stock concentration | Solvent | RT | m/z (+) |
|---|---|---|---|---|---|---|
| *Std. 1 – *ortho*-aryl-OH-6PPD-Q | 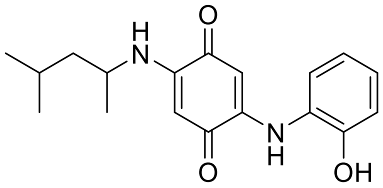 | C18H24N2O3 | 500 mg/L | DMSO:MeOH (1:1) | 21.40 | 315.1700 |
| *Std. 2 – *meta*-aryl-OH-6PPD-Q | 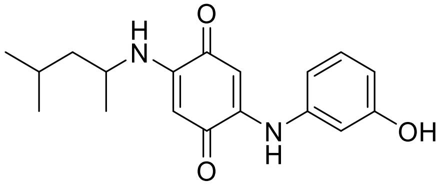 | C18H24N2O3 | 500 mg/L | DMSO:MeOH (1:1) | 21.03 | 315.1700 |
| *Std. 3 – *para*-aryl-OH-6PPD-Q | 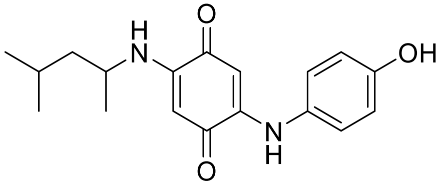 | C18H24N2O3 | 500 mg/L | DMSO:MeOH (1:1) | 20.64 | 315.1700 |
| *Std. 4 – $C_3$-alkyl-OH-6PPD-Q | 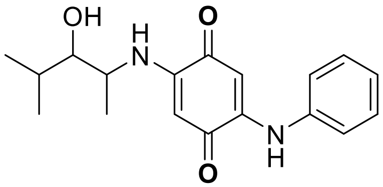 | C18H24N2O3 | 500 mg/L | DMSO:MeOH (1:1) | 20.24 | 315.1700 |
| *Std. 5 – $C_4$-alkyl-OH-6PPD-Q | 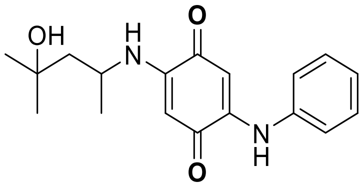 | C18H24N2O3 | 500 mg/L | DMSO:MeOH (1:1) | 19.48 | 315.1700 |

| | | | | | | |
|---|---|---|---|---|---|---|
| *Std. 6 – IPPD-Q | 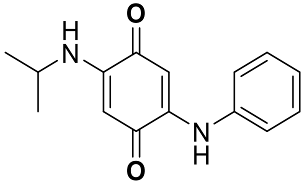 | C15H16N2O2 | 500 mg/L | DMSO:MeOH (1:1) | 19.68 | 257.1300 |
| *Std. 7 – CPPD-Q | 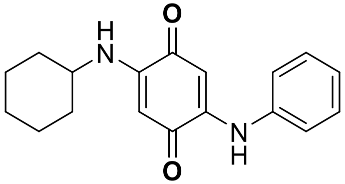 | C18H20N2O2 | 500 mg/L | DMSO:MeOH (1:1) | 21.59 | 297.1600 |
| *Std. 8 – HPPD-Q | 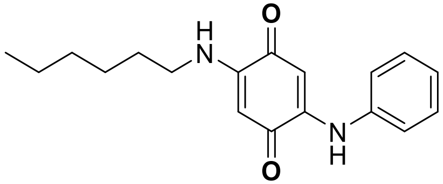 | C18H22N2O2 | 500 mg/L | DMSO:MeOH (1:1) | 22.07 | 301.1911 |
| *Std. 9 – OPPD-Q | 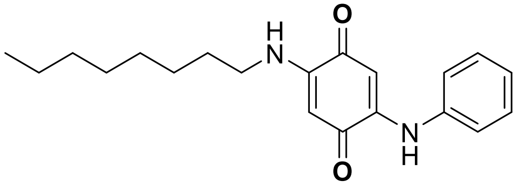 | C20H26N2O2 | 500 mg/L | DMSO:MeOH (1:1) | 23.09 | 327.2067 |
| *Std. 10 – DPPD-Q | 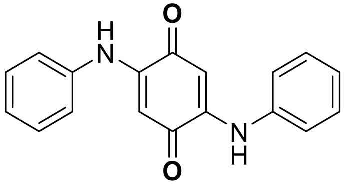 | C18H14N2O2 | 50 mg/L | DMSO | 20.77 | 291.1128 |
| *Std. 11 – DTPD-Q | 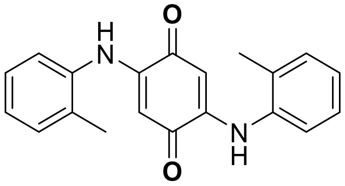 | C20H18N2O2 | 50 mg/L | DMSO | 21.41 | 319.1441 |

| | | | | | | |
|---|---|---|---|---|---|---|
| 6PPD-Q | 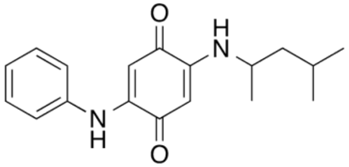 | C18H22N2O2 | 600 mg/L | DMSO | 5.37 (section S5.1)<br>21.87 (section S5.2) | 299 (section S5.1)<br>299.1754 (section S5.2) |
| 6PPD-Q d5 (IS) | 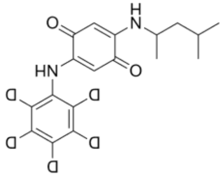 | C18H17D5N2O | 10 mg/L | MeOH | 21.84 | 304.2000 |

* Provided by Prof. Hui Peng, Department of Chemistry, and School of the Environment, Downtown Toronto (St. George)

**Supplementary Table S3.** Decrease in *para*-aryl-OH-6PPD-Q concentration in the cytotoxicity assay over the 24-h exposure period, calculated as $100 - (t_{24}/t_0 \times 100)$ for each biological replicate at a nominal concentration of 500 µg/L. * The nominal concentration in R1 of the no-cells control was 1000 µg/L.

| Cell line | R1 decrease (%) | R2 decrease (%) | R3 decrease (%) | Mean decrease (%) |
|---|---|---|---|---|
| RTBrain | 40.76 | 36.66 | 20.30 | 32.57± 12.57 |
| RTgill-W1 | 71.51 | 57.17 | 57.72 | 62.13 ± 8.12 |
| No-cells controls | 17.69* | 14.45 | 40.07 | - |

**Supplementary Table S4.** Standard curves used in the biotransformation experiments with 6PPD-Q and in the cytotoxicity assays with 6PPD-Q and BTPs, including calibration ranges and $R^2$ values. *Analysis marked with an asterisk was performed at EPFL; all other analyses were performed at Eawag.

| Cell line | Experiment & Replicate (R) number | Calibration curve | Calibration range (µg/L, online vial) | $R^2$ |
|---|---|---|---|---|
| RTBrain | Cytotoxicity assay – round 1* | 6PPD-Q | 50-0.1 | 0.99 |
| RTBrain | Biotransformation - R1 (48 h) | 6PPD-Q | 0.025-0.000390625 | 0.99 |
| RTBrain | Biotransformation - R2 (48 h) | 6PPD-Q | 0.025-0.000390625 | 0.99 |
| RTBrain | Biotransformation - R3 (48 h) | 6PPD-Q | 0.025-0.000390625 | 0.99 |
| RTBrain | Biotransformation - R1 (96 h) | 6PPD-Q | 0.025-0.000390625 | 0.99 |
| RTgill-W1 | Biotransformation - R1 (48 h) | 6PPD-Q | 0.025-0.000390625 | 0.99 |
| RTgill-W1 | Biotransformation - R2 (48 h) | 6PPD-Q | 0.025-0.000390625 | 0.99 |
| RTgill-W1 | Biotransformation - R3 (48 h) | 6PPD-Q | 0.025-0.000390625 | 0.99 |
| RTgill-W1 | Biotransformation - R1 (96 h) | 6PPD-Q | 0.025-0.00078125 | 0.99 |
| RTBrain/RTgill-W1 | Cytotoxicity assay round 2; BTP quantification biotransformation experiments | C3-alkyl-OH-6PPD-Q | 0.125-0.00390625 | 0.99 |
| RTBrain/RTgill-W1 | Cytotoxicity assay round 2; BTP quantification biotransformation experiments | C4-alkyl-OH-6PPD-Q | 0.125-0.00390625 | 0.98 |
| RTBrain/RTgill-W1 | Cytotoxicity assay round 2; BTP quantification biotransformation experiments | *meta*-aryl-OH-6PPD-Q | 0.0625-0.00390625 | 0.99 |
| RTBrain/RTgill-W1 | Cytotoxicity assay round 2; BTP quantification biotransformation experiments | *para*-aryl-OH-6PPD-Q | 0.0625-0.00390625 | 0.99 |
| RTBrain/RTgill-W1 | Cytotoxicity assay round 2; BTP quantification biotransformation experiments | *ortho*-aryl-OH-6PPD-Q | 0.0625-0.00390625 | 0.99 |
| RTBrain/RTgill-W1 | Cytotoxicity assay round 2 | IPPD-Q | 0.0625-0.00390625 | 0.99 |
| RTBrain/RTgill-W1 | Cytotoxicity assay round 2 | OPPD-Q | 0.25-0.015625 | 0.97 |
| RTBrain/RTgill-W1 | Cytotoxicity assay round 2 | DPPD-Q | 0.125-0.00390625 | 0.99 |
| RTBrain/RTgill-W2 | Cytotoxicity assay round 2 | CPPD-Q | 0.125-0.00390625 | 0.98 |
| RTBrain/RTgill-W3 | Cytotoxicity assay round 2 | HPPD-Q | 0.125-0.00390625 | 0.98 |
| RTBrain/RTgill-W4 | Cytotoxicity assay round 2 | DTPD-Q | 0.125-0.00390625 | 0.98 |

**Supplementary Table S5.** Positive control data from the cytotoxicity assays under the OECD TG 249. NE, not reliably estimated, indicates that the 95% confidence interval or Hill slope could not be estimated with sufficient precision.

| Positive control | AB EC50 | AB Hill slope | AB 95% CI | CFDA-AM EC50 | CFDA-AM Hill Slope | CFDA-AM 95% CI | NR EC50 | NR Hill slope | NR 95% CI | Chemical(s) tested | Cell line |
|---|---|---|---|---|---|---|---|---|---|---|---|
| (1) SDS | 4.90 | -1.88 to -1.33 | 4.35 to 5.51 | ~ 17.47 | NE | NE | 10.03 | -1.86 to -1.22 | 8.75 to 11.50 | *Ortho-*, *meta-*, *para-*aryl-OH-6PPD-Q | RTgill-W1 |
| (2) SDS | 3.80 | -1.39 to -0.98 | 3.26 to 4.44 | 12.88 | -9.96 to -1.69 | 9.33 to 17.79 | 6.857 | -2.63 to -1.85 | 6.30 to 7.46 | C3- and C4-alkyl-OH-6PPD-Q, IPPD-Q, HPPD-Q, OPPD-Q | RTgill-W1 |
| (3) SDS | 4.65 | -1.63 to -1.00 | 3.83 to 5.65 | 14.91 | -11.34 to -0.14 | 10.98 to 20.23 | 11.37 | -1.89 to -1.24 | 9.97 to 12.98 | CPPD-Q, *para*-aryl-OH-6PPD-Q, DPPD-Q, DTPD-Q; co-exposure 6PPD-Q & MK-571 | RTgill-W1 |
| (4) SDS | 4.20 | -1.80 to -0.77 | 3.01 to 5.88 | ~ 16.58 | NE | NE | 8.03 | -2.39 to -1.77 | 7.44 to 8.66 | *para*-aryl-OH-6PPD-Q | RTgill-W1 |
| (5a) SDS | 4.30 | -1.84 to -1.23 | 3.71 to 4.97 | ~ 16.91 | NE | NE | 7.37 | -6.25 to -3.18 | 7.11 to 7.64 | co-exposure 6PPD-Q & MK-571 | RTgill-W1 |
| (5b) SDS | 4.32 | -2.07 to -1.52 | 3.92 to 4.77 | 16.27 | -39.38 to 22.67 | 7.56 to 35.04 | 6.58 | -2.51 to -1.60 | 5.84 to 7.41 | | |
| (6) 3,4-DCA | 48.44 | -4.31 to -2.21 | 43.87 to 53.49 | 61.26 | -12.34 to 0.20 | 48.11 to 78.00 | ~ 54.37 | NE | NE | 6PPD-Q | RTBrain |

| | | | | | | | | | | | |
|---|---|---|---|---|---|---|---|---|---|---|---|
| (7) SDS | 3.84 | -1.51 to -0.83 | 2.95 to 5.00 | 23.44 | -2.17 to -0.44 | 14.63 to 37.56 | 27.94 | -16.30 to 11.01 | 4.92 to 158.60 | *Ortho-*, *meta-*, *para-*aryl-OH-6PPD-Q | RTBrain |
| (8) SDS | 5.89 | -3.89 to -2.97 | 5.69 to 6.11 | 15.39 | -2.84 to -1.09 | 12.20 to 19.42 | 11.88 | -0.99 to -0.05 | 5.55 to 25.45 | C3- and C4-alkyl-OH-6PPD-Q, IPPD-Q, CPPD-Q, HPPD-Q, OPPD-Q | RTBrain |
| (9a) SDS | 3.86 | -1.74 to -1.06 | 3.19 to 4.68 | 10.74 | -3.92 to -2.17 | 9.54 to 12.09 | 9.22 | -2.96 to -2.06 | 8.56 to 9.92 | DPPD-Q, DTPD-Q, *para*-aryl-OH-6PPD-Q | RTBrain |
| (9b) SDS | 4.70 | -1.52 to -0.75 | 3.44 to 6.43 | 15.43 | -10.57 to -1.58 | 12.64 to 18.84 | 7.84 | -2.43 to -1.27 | 6.54 to 9.39 | | |
| (10) SDS* | 3.62 | -2.33 to -1.50 | 3.19 to 4.10 | 10.92 | -8.96 to -0.31 | 7.83 to 15.22 | 5.36 | -3.97 to -0.78 | 3.79 to 7.58 | *para*-aryl-OH-6PPD-Q | RTBrain |

** Only one control well was included in the analysis due to cell detachment in the remaining wells.*

**Supplementary Table S6.** Summary of the acceptance ranges for 3,4-DCA and SDS in RTgill-W1 cells, based on previously established ranges, and the SDS acceptance ranges calculated for RTBrain cells in this study as the mean ± 2.5 SD.

| Cell line | Positive Control | AB range | CFDA-AM range | NR range | Reference |
|---|---|---|---|---|---|
| RTgill-W1 | 3,4 DCA | 43.6 mg/L ± 6.1 mg/L (2½ SD range: 28.4 to 58.9 mg/L) | 62.5 mg/L ± 18.9 mg/L (2½ SD range: 15.3 to 109.8 mg/L) | 58.6 mg/L ± 18.6 mg/L (2½ SD range: 12.1 to 105.0 mg/L) | (78) |
| RTgill-W1 | SDS | 4.27 mg/L ± 1.037 (2½ SD range: 1.68 to 6.86 mg/L) | 9.9 mg/L ± 2.131 (2½ SD range: 4.57 to 15.23 mg/L) | 6.7 mg/L ± 1.523 (2½ SD range: 2.89 to 10.51 mg/L) | (*82*) |
| RTBrain | SDS | 4.3 ± 1.05 mg/L (2½ SD range: 1.7–6.9 mg/L) | 16.6 ± 4.56 mg/L (2½ SD range: 5.2–28.0 mg/L), | 9.0 ± 2.59 mg/L (2½ SD range: 2.5–15.5 mg/L) | This study |

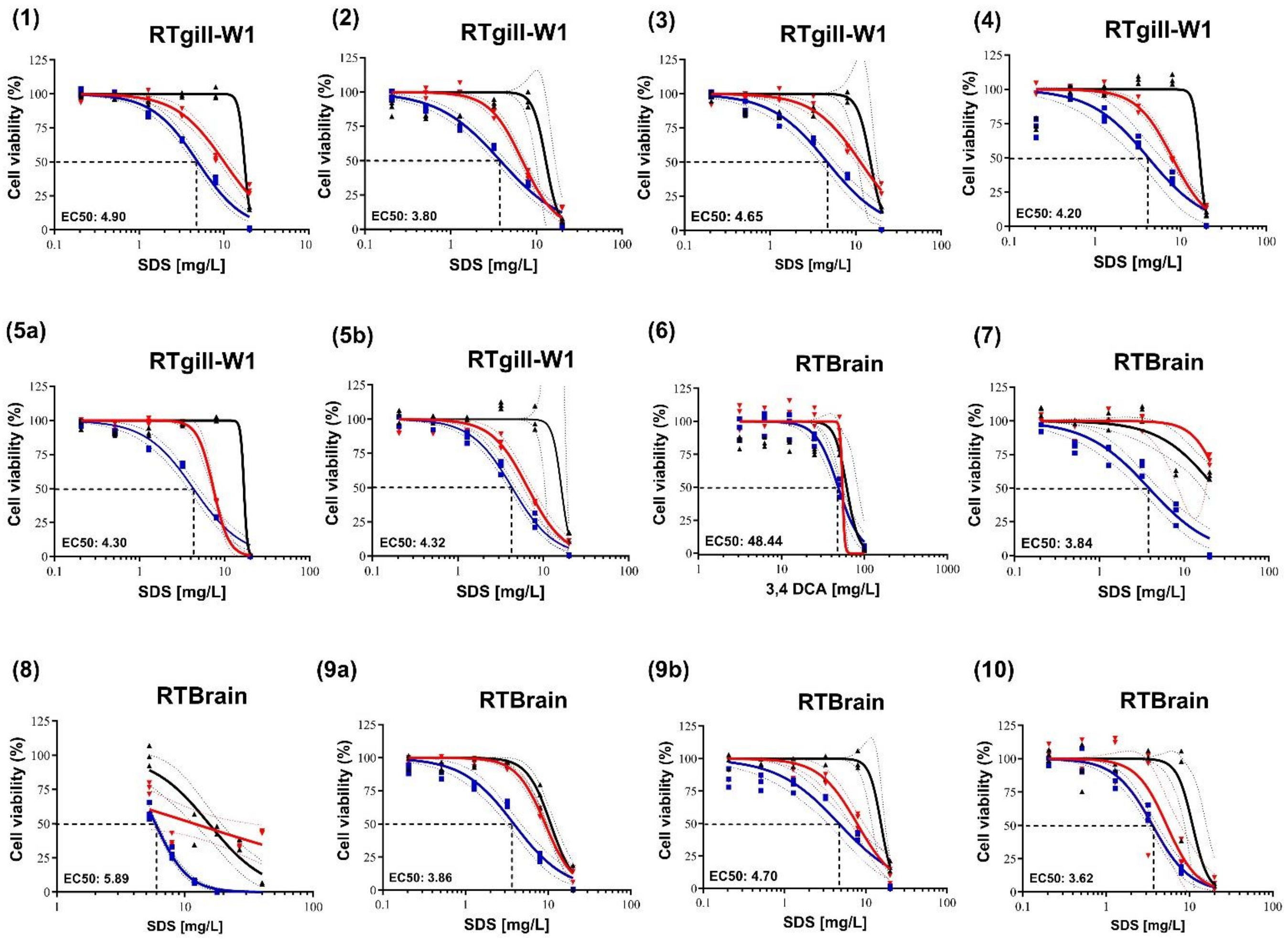

**Supplementary Figure S16.** Positive control data from cytotoxicity assays conducted according to OECD TG 249. Numbers correspond to the entries in Supplementary Table S5. Dose-response curves represent cell metabolic activity (alamarBlue ™; blue squares), cell membrane integrity (CFDA-AM; black triangles) and lysosomal integrity (Neutral Red; red inverted triangles) after 24 h of exposure. Data are expressed as percentage of solvent control (0.5% (v/v) DMSO)
